# A Fungal Natural Product that Inhibits Plant Cellulose Biosynthesis by disrupting Cellulose Synthase Complexes

**DOI:** 10.64898/2026.04.12.718067

**Authors:** Zhongshou Wu, Lu Liu, Wenyu Han, Xingbo Cai, Pixian Xiao, Zuodong Sun, Chunsheng Yan, Silvana Reid, Yun Chen, Zhonghua Ma, Yi Tang, Steven E. Jacobsen

**Author notes:** These authors contributed equally to this work (Zhongshou Wu, Lu Liu, Wenyu Han). **Author Contributions:** Z.W., L.L., Y.T. and S.E.J. designed the research, interpreted data, and wrote the manuscript; Z.W. performed most experiments. L.L. performed the NGS analysis, confocal analysis and generated multi-drugs resistant mutants and wrote the manuscript. W.H. and Z.S. isolated MDD and related compounds. X.C. and P.X. performed some confocal analysis. Y.C. and Z.M. analyzed the data. **Competing Interest Statement:** Authors declare that they have no competing interests.

## Abstract

Cellulose, a primary component of plant cell walls, is synthesized by cellulose synthase complexes (CSCs) at the plasma membrane. Targeting this process with cellulose biosynthesis inhibitors (CBIs) has significantly advanced our understanding of plant cell wall formation and provided valuable compounds for herbicide development. Here, we identified a fungal natural product, 8-methyldichlorodiaporthin (MDD), as a broad-spectrum plant CBI. Structure-activity relationship analyses demonstrate that methylation modifications on the isocoumarin ring and chlorination of side chain are crucial for MDD-induced growth inhibition. A chemical forward genetic screen in *Arabidopsis thaliana* revealed two semi-dominant CESA1 mutations, causing A903T and H1024Y substitutions, that confer insensitivity to MDD. Both mutations locate to transmembrane domains of CESA1, and we show that MDD depletes CSCs from the plasma membrane and reduces cellulose content. Further genetic analyses indicate that the *cesa1*^*mddi1-1*^ *A903T* mutant also confers resistance to CBIs quinoxyphen and C17, but not to CBIs isoxaben, indaziflam, or ES20. Stacking additional point mutations conferring resistance to other CBIs, *cesa3*^*ixr1-1*^ *G998D*, and *cesa6*^*es20-r3*^ *G935E* into the *cesa1*^*mddi1-1*^ *A903T* background yields multiple-drugs resistant lines that maintain normal growth. These findings establish MDD, as a novel, natural CBI that likely targets CESA1, thereby extending our understanding of CSC regulation and abilities to develop multi-drugs resistant crop varieties. These findings offer new perspectives for weed management and plant biotechnology.

**Significance Statement:** Cellulose, a fundamental structural component of plant cell walls, is synthesized by cellulose synthase complexes (CSCs) and represents a critical herbicide target. While synthetic cellulose biosynthesis inhibitors (CBIs) like isoxaben and quinoxyphen have helped in the elucidation of CSC function and aided in weed control, natural CBIs remain largely undiscovered. Here, we identify 8-methyldichlorodiaporthin (MDD), a fungal-derived isocoumarin natural product, as a CBI that inhibits plant growth by depleting CSCs from the plasma membrane. Genetic screens reveal MDD-resistant *cesa1* mutations, and combining these with other CBI-resistant alleles yields multi-herbicide resistant plants that can grow normally. This research enhances our understanding of cellulose biosynthesis and paves the way for multi-herbicide resistant crops with agricultural benefits.

## Introduction

Cellulose microfibrils, crystalline polymers of β-1,4-D-glucose, play a vital role in providing structural support and rigidity to plant cell walls. Within the plasma membrane, multiple monomeric cellulose synthases (CESAs) form a larger rosette-like structure known as the cellulose synthase complex (CSC) and are responsible for cellulose production (1-4). In *Arabidopsis*, the CSC that synthesizes cellulose in the primary cell wall consists of CESA1, CESA3, and either CESA6 or a CESA6-like subunit (CESA2, CESA5, or CESA9), whereas the CSC in the secondary cell wall is composed of CESA4, CESA7, and CESA8 (5-8).

Cellulose biosynthesis inhibitors (CBIs) are chemical compounds that disrupt the synthesis of cellulose in plants, thereby causing cell swelling, affecting CSC subcellular localization, and impacting plant growth and development (9-11). CBIs include both synthetic compounds—such as isoxaben, quinoxyphen, C17, flupoxam and triazofenamide—and a small number of natural products (e.g., thaxtomin A). Different CBIs act by distinct mechanisms: some (isoxaben, quinoxyphen, C17 and related compounds) cause removal or destabilization of CSCs from the plasma membrane, Endosidin20 (ES20) targets the catalytic site of CESA6, and other agents (e.g., indaziflam, morlin) influence CSC abundance or trafficking indirectly (9, 12-15). Because these inhibitors perturb a fundamental process in cell expansion, they are valuable both as herbicides and as tools to dissect CSC structure–function relationships. Genetic screens have shown that single amino-acid substitutions in CESA proteins can confer resistance to specific CBIs, and many resistance mutations cluster in transmembrane domains (9, 14, 16-19), highlighting likely functional or binding hotspots. All of these compounds have also been demonstrated to have inhibitory effects on a range of plant species, with isoxaben being widely used as an herbicide to control broadleaf weeds (6, 13, 14, 16-18, 20-23). Notably, mutants that confer resistance to specific CBIs do not typically exhibit cross-resistance to other types (14, 20, 24-26). The diversity of CBIs and the mutation-resistance data emphasize the value of identifying new inhibitors and their targets to expand herbicide modes of action and to probe CSC biology.

Microbial natural products remain an underexplored source of CBIs. In microbial genomes, biosynthetic genes are typically found in biosynthetic gene clusters (BGCs). Advances in genomics and metabolomics have enabled the discovery of new compounds and their biological targets (27-29). For example, expressing a relatively well-conserved fungal BGC found in diverse fungi in the heterologous *Aspergillus nidulans* host led to the successful production of isocoumarin compounds 8-methyldichlorodiaporthin (MDD) and dichlorodiaporthin (DD) (30). These are polyketide derived compounds that have undergone significant modifications, including *gem-*dichlorination, methylation and C-C bond cleavage. While these compounds were previously isolated and rediscovered through genome mining efforts (30, 31), the biological activity and target remained elusive. Synthetic isocoumarin compounds have been shown to be potential herbicides (32). Given the ecological niche of fungi in plant-fungal interactions, we hypothesized that MDD and/or DD could have potential roles in inhibiting essential plant enzymes and could serve as herbicides.

Here, we discovered that MDD is a broad-spectrum plant growth inhibitor. Structure-activity relationship analysis revealed the critical roles of aforementioned chemical features, including both 6-OMe and 8-OMe groups, and the *gem*-dichloro group in effective root growth inhibition. A forward chemical genetic screen in *Arabidopsis thaliana* revealed that substitutions in CESA1 (*cesa1*^*mddi1-1*^ *A903T* and *cesa1*^*mddi1-2*^ *H1024Y*) confer resistance to MDD. MDD inhibits cellulose biosynthesis and depletes CSCs from the plasma membrane. The *cesa1*^*mddi1-1*^ *A903T* mutant is resistant to two additional CBIs, quinoxyphen and C17. Additionally, we have successfully developed mutants with resistance to four CBIs by combining *cesa1*^*mddi1-1*^ *A903T* with *cesa3*^*ixr1-1*^ *G998D* or *cesa6*^*ixr2-1*^*R1064W* or *cesa6* ^*es20-r3*^ *G935E*. Moreover, a triple mutant combining *cesa1*^*mddi1-1*^ *A903T, cesa3*^*ixr1-1*^ *G998D* and *cesa6* ^*es20-r3*^ *G935E* shows resistance to five CBIs, MDD, quinoxyphen, C17, isoxaben and ES20. The bioactivity study of MDD not only expands herbicide options for weed control but also helps mitigate herbicide-driven resistance evolution in grasses by enabling the rotation or combination of herbicides with alternative modes of action.

## Results

### MDD is a broad-spectrum plant growth inhibitor

The herbicidal activity of MDD (Figure 1A) was discovered through an *Arabidopsis*-based screening of a collection of natural fungal compounds obtained from our genome mining efforts. MDD, a known natural product, was rediscovered through biosynthesis in *A. nidulans* by heterologous reconstitution of the *dia* BGC originally found in *Aspergillus oryzae* (30). The defining structural features of MDD include an isocoumarin core derived from a nonreducing polyketide synthase; a *gem*-dichloro group derived from a flavin-dependent halogenase and two methoxy groups at C-6 and C-8 from SAM-dependent *O-*methyltransferase.

**Fig. 1.**
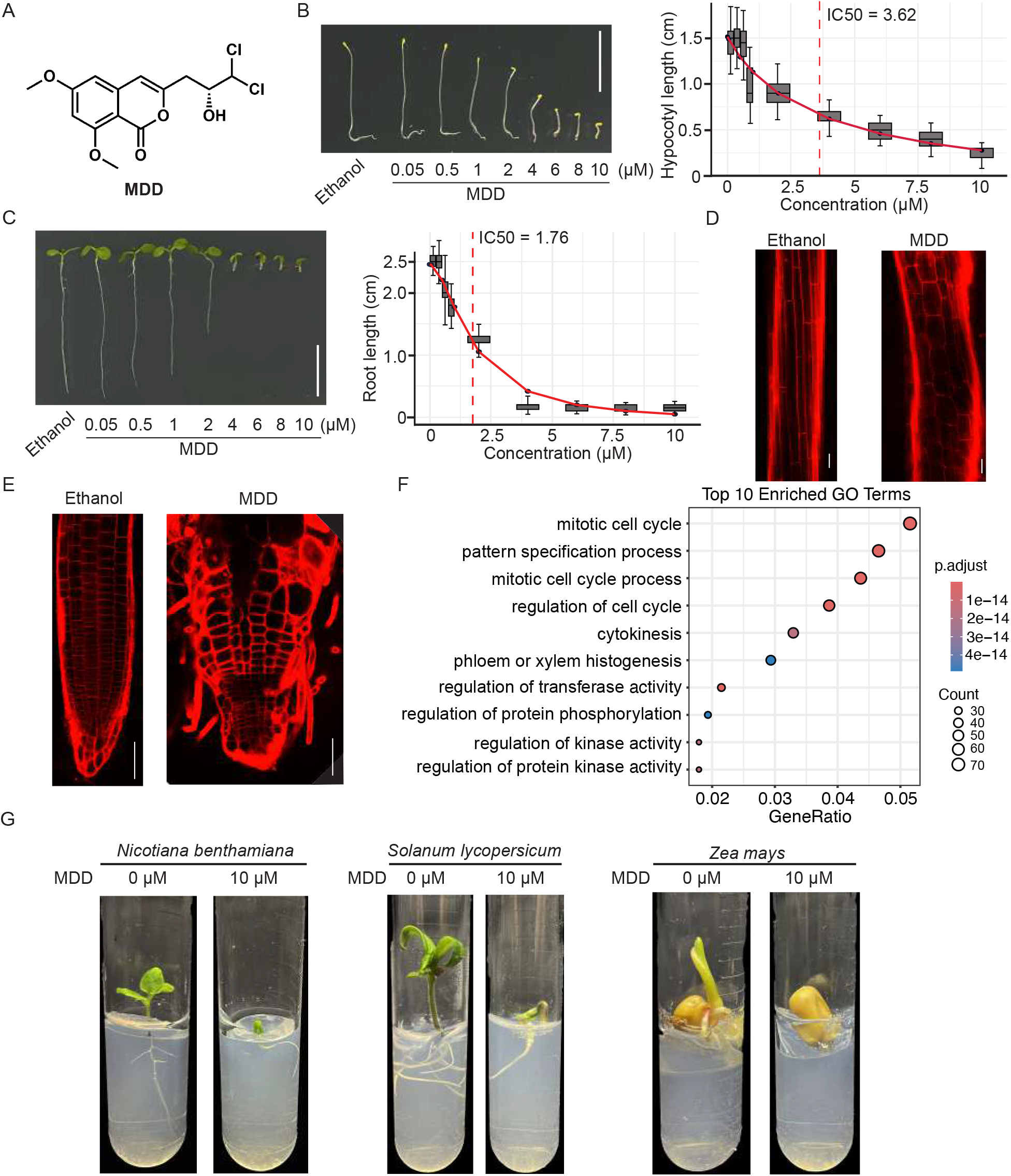
MDD is a broad-spectrum plant growth inhibitor. A.Chemical structure of MDD. B.Representative image and quantification of hypocotyl growth of 5-day-old dark-grown Arabidopsis seedlings grown on ½ MS medium supplemented with Ethanol or increasing concentrations of MDD. Bars = 1.0 cm. C.Representative image and quantification of root growth of 7-day-old light-grown Arabidopsis seedlings grown on ½ MS medium supplemented with Ethanol or increasing concentrations of MDD. Bars = 1.0 cm. D.Representative image of hypocotyl cells of 4-day-old dark-grown Arabidopsis with Ethanol or 10 μM MDD treatment. Bars = 50 μm E.Representative image of root cells of 4-day-old light-grown Arabidopsis with Ethanol or 10 μM MDD treatment. Bars = 50 μm. F.Top 10 enriched GO terms of downregulated genes of MDD treated WT plants. G.Representative image of 14-day-old light-grown *Nicotiana Benthamiana, Solanum lycopersicum* and *Zea mays*.

The addition of MDD resulted in significant inhibition of *Arabidopsis* wild-type (WT) Col-0 hypocotyl and root growth in a dose-dependent manner (Figure 1B-C and Fig. S1A-B). The concentration of MDD that caused a 50% reduction in hypocotyl and root growth (IC50) was determined to be 3.62 μM and 1.76 μM, respectively (Figure 1B-C). When exposed to MDD, hypocotyl and root cells exhibited severe swelling (Figure 1D-E and Fig. S1C-D), indicated by a notable decrease in cell length and an increase in cell width in both hypocotyls and the root elongation zone (Figure 1D-E and Fig. S1C-D). Transcriptomic profiling revealed that genes downregulated by MDD were significantly enriched in core cell cycle processes, particularly mitotic cell cycle and cytokinesis (Figure 1F). Consistent with this, MDD⍰treated roots also exhibited a reduced number of meristematic cells compared with ethanol⍰treated controls (Fig. S1E). Additionally, MDD significantly inhibited the growth of various dicotyledonous and monocotyledonous plant species, including tobacco, tomato, and maize (Figure 1G), demonstrating that MDD is an effective broad-spectrum plant growth inhibitor.

To further investigate the structure-activity relationship of MDD-induced plant growth inhibition, *Arabidopsis* plants were treated with MDD and its structural analogues (Figure 2A). These MDD analogues (DD and MA2-MA9) are either on-pathway intermediates or shunt products, isolated during the heterologous biosynthesis experiments (30). Notably DD is the C-8 desmethyl version of MDD and is also a known natural product from previous isolation (30). Formation of DD instead of MDD is attributed to the lack of one methyltransferase in the corresponding BGCs compared to that of MDD (30). Based on the structure-activity relationship studies, only DD and MA-6 caused significant inhibition, reducing WT root growth by 40% and 50% respectively (Figure 2B-C). The variations in root length among plants treated with MDD, DD, and MA-2 highlight the critical roles of the 6-OMe and 8-OMe groups in effective root growth inhibition. Additionally, the unique *gem-* dichloro moiety on the isocoumarin C-3 alkyl chain plays a pivotal role in the observed biological activity, since nonchlorinated analogues MA-5, MA-6 and MA-7 exhibited weaker or negligible inhibitory effect.

**Fig. 2.**
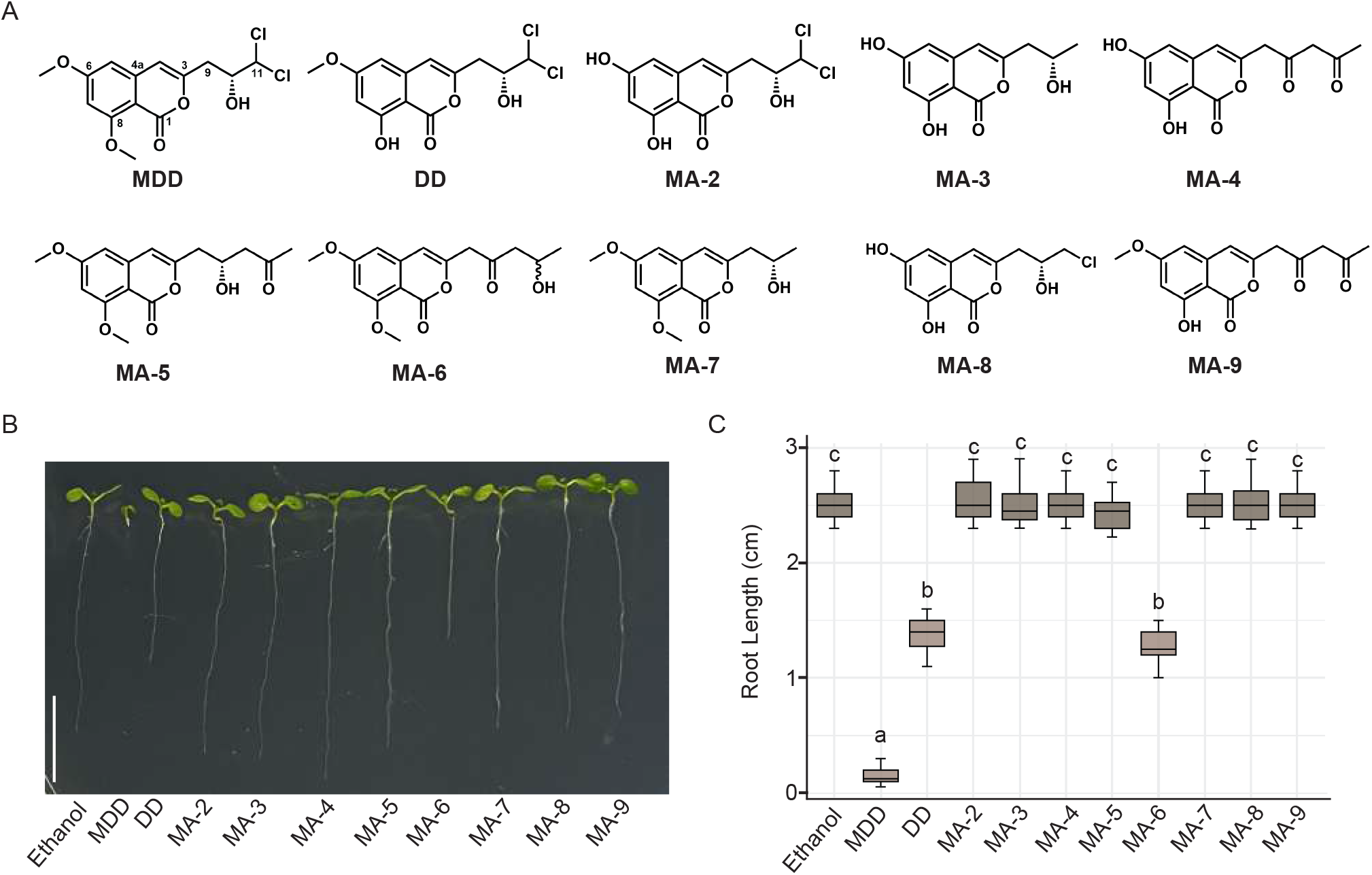
Methylation patterns on the isocoumarin ring and chlorination of side chain are crucial for MDD-induced growth inhibition. A.Chemical structures of MDD, DD and 8 analogs (MA-2-9). B.Representative image of 7-day-old Arabidopsis light-grown seedlings grown on ½ MS medium supplemented with Ethanol or 10 μM of MDD and indicated analogs. Bars = 1.0 cm. C.Quantification of root growth of 7-day-old light-grown Arabidopsis seedlings grown on ½ MS medium supplemented with Ethanol or 10 μM of MDD and indicated analogs. Statistical significance is indicated by different letters (*p*□<□0.01). Error bars represent means□±□standard deviations (n□=□3).

### Resistance to MDD is conferred by semi-dominant CESA1 alleles

To elucidate the mode of action of MDD in plant inhibition, a chemical forward genetic screen was conducted to identify mutants that are insensitive to MDD. Approximately 5,000 *Arabidopsis* WT seeds were mutagenized using ethyl methanesulfonate (EMS), and approximately 30,000 M2 seeds were screened for growth insensitivity to MDD. *mddi1-1* and *mddi1-2* (*<u>MDD</u>-insensitive*) mutants were isolated, both exhibiting loss of sensitivity to MDD (Figure 3A-B and Fig. S1A-B).

**Fig. 3.**
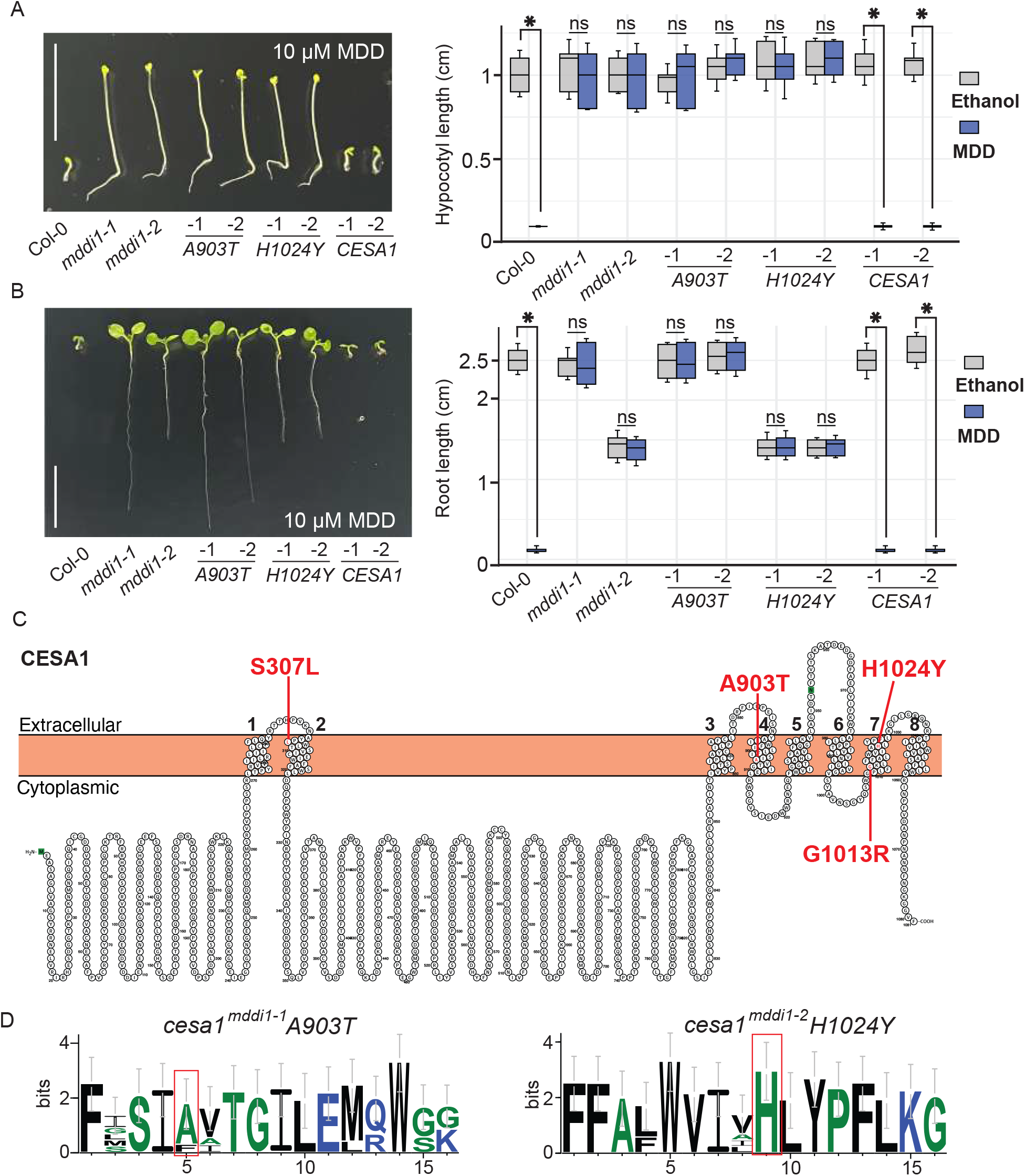
Mutations in CESA1 cause reduced sensitivity to MDD. A.Representative image and quantification of hypocotyl growth of 5-day-old dark-grown Arabidopsis seedlings grown on ½ MS medium supplemented with 10 μM MDD. Bars = 1.0 cm. Statistical significance is indicated by an asterisk (*p*□<□0.01). Error bars represent means□±□standard deviations (n□=□3). B.Representative image and quantification of root growth of 7-day-old light-grown Arabidopsis seedlings grown on ½ MS medium supplemented with 10 μM MDD. Bars = 1.0 cm. Statistical significance is indicated by asterisks (*p*□<□0.01). Error bars represent means□±□standard deviations (n□=□3). C.Predicted topology of CESA1. □ocations of the mutated amino acids that cause reduced sensitivity to MDD are labeled with red color. D.Sequence logo assessment of residues in the *mddi1-1, mddi1-2* mutation regions of primary cell wall CESA proteins illustrates the location and conservation of the mutated alanine and histidine residues in CESA1.

To map the *mddi* mutants, homozygous *mddi* mutants were backcrossed with WT to create a mapping population. Sixty F2 plants that were insensitive to MDD were selected for Illumina sequencing. Single nucleotide polymorphism (SNP) frequency analysis identified a single causal region for both *mddi* mutants located at the end of chromosome 4 (Fig. S2A). Detailed analysis of mutations in each *mddi* mutant revealed that *CESA1* (*AT4G32410*) was the only gene with mutations in both mutants. The *mddi1-1* mutant carries a G-to-A mutation at nucleotide 2707, resulting in an Ala903Thr substitution in the fourth transmembrane domain (Figure 3C). The *mddi1-2* mutant has a C-to-T change at nucleotide 3070, causing a His1024Tyr substitution in the seventh transmembrane domain (Figure 3C). Among primary cell wall CESAs (CESA1, 3, 6, 2, 5, 9), the mutated histidine at position 1024 is absolutely conserved, while alanine at position 903 is highly conserved. Additionally, both residues are flanked by highly conserved residues (Figure 3D).

When *mddi1-1* and *mddi1-2* were backcrossed with WT, the resulting F1 plants were partially resistant to MDD, indicating that the *mddi* mutations are semi-dominant (Fig. S2B). To confirm that mutations in CESA1 are responsible for the MDD-insensitive phenotype, we performed a transgene complementation analysis. We introduced N-terminal YFP-tagged WT CESA1 and CESA1 carrying the A903T or H1024Y substitutions into heterozygous *cesa1* knockout mutants (*SAIL_278_E08*). Introducing the A903T or H1024Y substitutions fully rescued the lethality of the *cesa1* null mutant (Fig. S2C) and conferred insensitivity to MDD (Figure 3A–B), whereas introduction of wild-type CESA1 rescued lethality but failed to generate MDD insensitivity (Figure 3A–B and Fig. S2C).These findings indicate that the missense mutations in CESA1 are sufficient to confer tolerance to MDD.

### MDD depletes CSCs from the plasma membrane and inhibits cellulose biosynthesis

CESA1 encodes a cellulose synthase isomer, which colocalizes with CESA3 and CESA6 to the plasma membrane and forms CSCs involved in microfibril production (1). We examined the localization and dynamics of CSCs under normal conditions and following MDD treatment. In ethanol control treated hypocotyl epidermal cells, GFP-CESA3 was prominently associated with the plasma membrane. However, when treated with MDD for 5 hrs, the GFP-CESA3 signals at the plasma membrane were significantly reduced (Figure 4A-B), suggesting that MDD depletes CSC complexes from the plasma membrane. Consistent with this observation, MDD treatment decreased the crystalline cellulose content in light-grown seedlings (Figure 4C). Both *cesa1*^*mddi1-1*^ *A903T* and *cesa1*^*mddi1-2*^ *H1024Y* mutants exhibited slightly lower cellulose content as compared to WT Col-0, but were insensitive to MDD inhibition (Figure 4C). Additionally, treatments with MDD greatly reduced the velocity of labeled CSC complexes (Figure 4D). These results collectively indicate that MDD depletes CSCs from the plasma membrane and inhibits their role in cellulose biosynthesis.

**Fig. 4.**
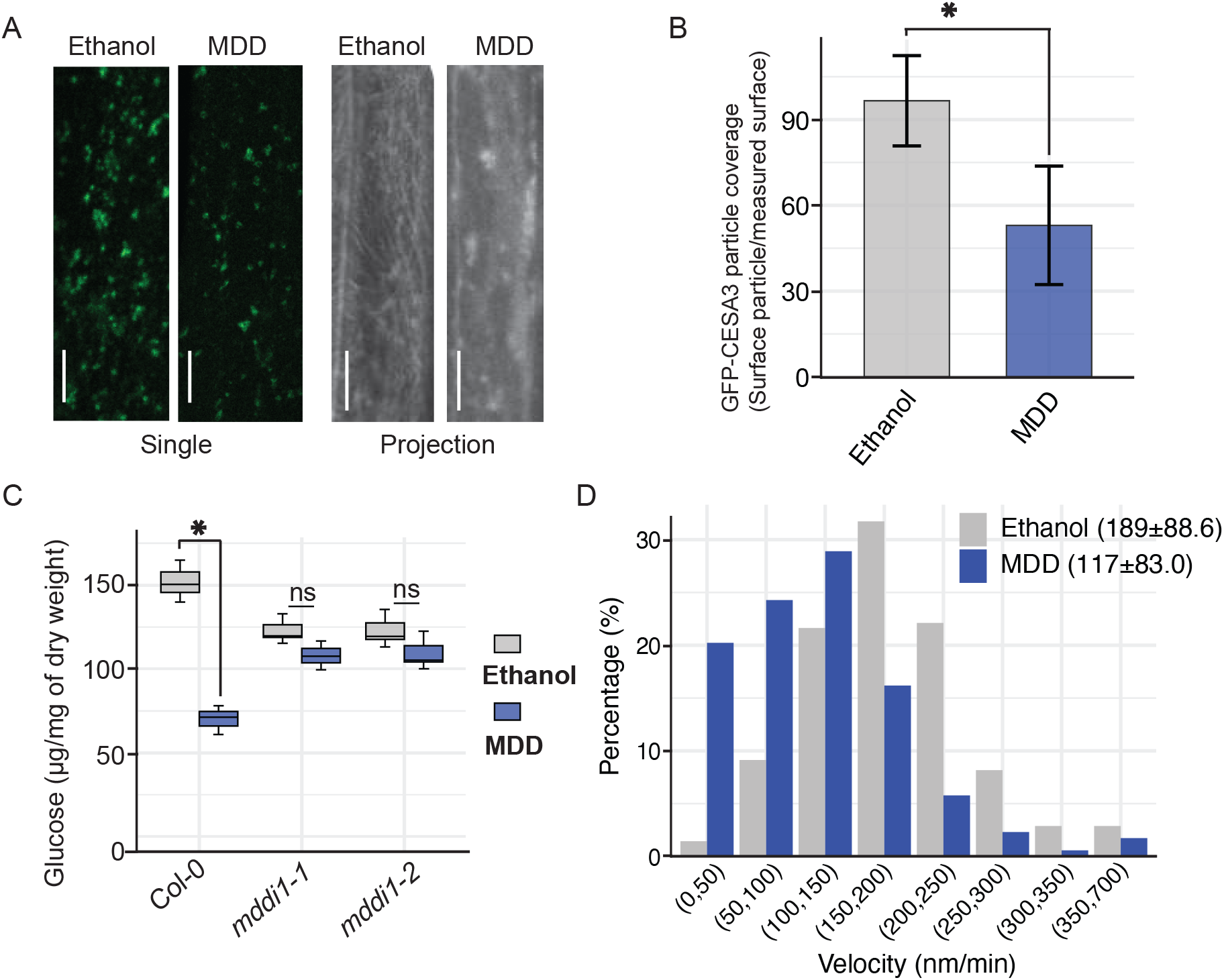
MDD depletes CSCs from the plasma membrane and inhibits cellulose biosynthesis. A.Representative single frame and time-projected images (4 min 24 frames) of CESA3-GFP in 5 hrs Ethanol- or 100 μM MDD-treated hypocotyl cells. Scale bar = 10 μm. B.Quantification of plasma membrane-localized GFP-CESA3 in hypocotyl cells after ethanol or 100 μM MDD treatment for 5 hrs. Error bars represent means□±□standard deviations (n□=□80 cells in each treatment). C.Glucose content of 7-day-old light-grown Arabidopsis seedlings without and with 10 μM MDD. Statistical significance is indicated by asterisk (*p*□<□0.01). Error bars represent means□±□standard deviations (n□=□3). D.Histogram with the frequencies of GFP-CESA3 particle velocity in hypocotyl cells after ethanol or 100 μM MDD treatment for 5 hrs. Data in the chart represent mean□±□standard deviation.

### MDD sensitivity remains unaltered by CSC complex quantity or regulatory mechanisms at the plasma membrane

CSCs are assembled in the Golgi and subsequently transported to the plasma membrane with the assistance of vesicle trafficking proteins and the trans-Golgi network (33). CESA INTERACTIVE PROTEIN 1 (CSI1), also known as POM-POM2 (POM2), links the CSC to cortical microtubules, guiding the movement of the CSC (34, 35). PATROL1 (PTL1) interacts with CSI1/POM2 and exocyst subunits to facilitate the delivery of CSC vesicles to the plasma membrane (36). In contrast, the SHOU4 protein regulates cellulose synthesis by limiting CSC exocytosis, and increased abundance of CESA proteins were observed at the plasma membrane in the *shou4* mutant background (37). To investigate whether MDD impacts these CSC-mis-accumulation mutants at the plasma membrane, we planted the *csi1-3* (*SALK_138584*), *patrol1-2* (*SALK_018676C*), and *shou4-3* (*GK793F10*) T-DNA insertion lines on ½ MS plate with MDD. However, none of these mutants were resistant to MDD (Fig. S3A-B), suggesting that MDD may also affect the functionality or operational efficiency of the CSCs, despite its pronounced effect on their abundance at the plasma membrane.

### MDD inhibits cellulose biosynthesis using a similar mode of action as quinoxyphen and C17 but operates differently than isoxaben, indaziflam, and ES20

The chemical structures of CBIs, including isoxaben, indaziflam, C17, and ES20 are distinct (Figure 1A and Figure 5A), yet all these compounds can inhibit cellulose biosynthesis (12, 13, 19, 20). To explore whether other known CBIs share a similar mechanism of action, we examined the toxicity of MDD on *Arabidopsis* mutants resistant to established CBIs. Isoxaben-resistant mutants *cesa3*^*ixr1-1*^ *G998D, cesa3*^*ixr1-2*^ *T942I, cesa6*^*ixr2-1*^ *R1064W* contain substitutions in the C-terminal transmembrane regions of the corresponding CESAs, where the *mddi* mutations are found. However, none of these mutants showed resistance to MDD (Fig. S4A). ES20 inhibits cellulose biosynthesis by targeting the catalytic domain of cellulose synthase, and most mutations that reduce sensitivity to ES20 are located in the central cytoplasmic domain. None of the tested ES20-resistant (*es20r*) mutants exhibited resistance to MDD (Fig. S4B). Interestingly, two C17-resistant mutants (*cesa1*^9R^ *G1013R* and *cesa1*^*18A1*^ *S307L*) showed partial insensitivity to MDD (Fig. S4C). The mutated glycine at position 1013 and serine at 307 are absolutely conserved and flanked by several highly conserved residues (Fig. S4D).

**Fig. 5.**
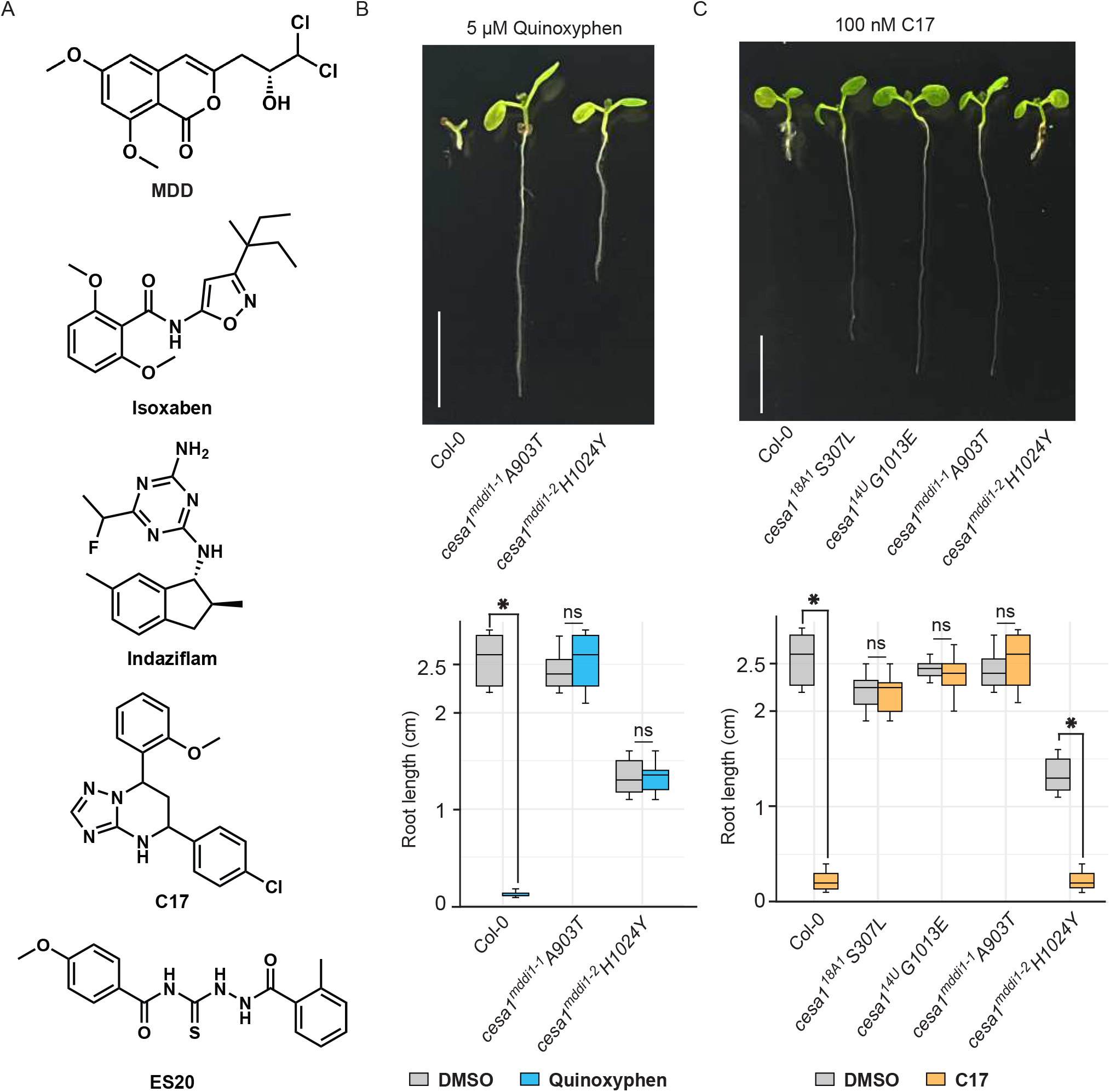
MDD inhibits cellulose biosynthesis using the same mode of action as quinoxyphen and C17. A.Chemical structures of MDD, isoxaben, indaziflam, C17 and ES20. B.Representative image and quantification of Col-0, *cesa1*^*mddi1-1*^ *A903T* and *cesa1*^*mddi1-2*^ *H1024Y* root growth of 7-day-old light-grown Arabidopsis seedlings grown on ½ MS medium supplemented with 5 μM Quinoxyphen. Bars = 1.0 cm. C.Representative image and quantification of Col-0, *cesa1*^*18A1*^ *S307□, cesa1*^*14U*^ *G1013E, cesa1*^*mddi1-1*^ *A903T* and *cesa1*^*mddi1-2*^ *H1024Y* root growth of 7-day-old light-grown Arabidopsis seedlings grown on ½ MS medium supplemented with 100 nM C17. Bars = 1.0 cm.

We also tested whether the *mddi* mutants exhibit altered sensitivity to other CBIs such as isoxaben, quinoxyphen, C17, and ES20. When treated with 10 nM isoxaben, or 1 μM ES20, the *mddi* mutants were inhibited to a similar level as the WT plants (Fig. S5A-B). In contrast, the *cesa1*^*mddi1-1*^ *A903T* mutant showed complete resistance to 5 μM quinoxyphen and 100 nM C17, while the *cesa1*^*mddi1-2*^ *H1024Y* mutant was completely resistant to 5 μM quinoxyphen but remained susceptible to 100 nM C17 (Figure 5B-C). Notably, *cesa1*^*mddi1-1*^ *A903T* contains a substitution at the same residue as *cesa1*^*aegeus*^ *A903V*, which is the only reported quinoxyphen-insensitive mutant (18). The insensitivity of *cesa1*^*mddi1-1*^ *A903T* to quinoxyphen and C17 also suggest that these two CBIs may function by a similar mode of action. Indeed, C17 insensitive mutants (*cesa1*^*9Q*^ *L872F, cesa1*^*14U*^ *G1013E, cesa1*^*9R*^ *G1013R* and *cesa1*^*18A1*^ *S307L*) showed partial resistant to quinoxyphen (Fig. S6A-B). The differing sensitivities of the *mddi* mutants to these CBIs suggest that MDD may share common characteristics in affecting cellulose synthesis with quinoxyphen and C17, but not with isoxaben, indaziflam, or ES20.

### Generation of multiple CBIs-resistant mutants

The *cesa1*^*mddi1-1*^ *A903T* mutant exhibited resistance to multiple CBIs, including quinoxyphen, C17, and MDD (Figure 3A-B and Figure 5B-C). To attempt to further broaden resistance, we stacked the isoxaben-resistant substitutions *cesa3*^*ixr1-1*^ *G998D* or *cesa6*^*ixr2-1*^ *R1064W* with *cesa1*^*mddi1-1*^ *A903T*. The resulting double mutants, *cesa1*^*mddi1-1*^ *A903T cesa3*^*ixr1-1*^ *G998D* or *cesa1*^*mddi1-1*^ *A903T cesa6*^*ixr2-1*^ *R1064W*, were able to tolerate combination treatments that included 10 μM MDD, 10 nM isoxaben, 5 μM quinoxyphen, and 50 nM C17 (Fig. S7A-B). When we reduced the concentrations of the inhibitors by 75%, the single mutants became sensitive to the combination, while the double mutants remained fully resistant (Fig. S7A-B). We next introduced the ES20-resistant mutant *cesa6* ^*es20-r3*^ *G935E* with *cesa1*^*mddi1-1*^ *A903T*. This double mutant likewise resisted a mixture of 10 μM MDD, 1 μM ES20, 5 μM quinoxyphen, and 50 nM C17 (Fig. S8A-B). Remarkably, a triple mutant carrying *cesa1*^*mddi1-1*^ *A903T, cesa3*^*ixr1-1*^ *G998D* and *cesa6* ^*es20-r3*^ *G935E* achieved resistance to five CBIs simultaneously, although slight growth inhibition was observed (Figure 6A-B). We attributed that minor inhibition to solvent effects arising from the relatively high total solvent volume used when applying multiple inhibitors. To address this, we reduced all herbicide concentrations to one-fifth of the original application levels. Under these lower⍰concentration conditions, the triple mutant showed clear resistance to all five CBIs, whereas single mutants remained susceptible (Figure 6A-B). At these reduced concentrations, solvent alone did not cause observable growth inhibition (Figure 6A-B). Although *cesa* null mutants were not viable, all CBI-resistant single and multiple mutants displayed wild-type morphology under our growth conditions (Figure 6C-D), demonstrating that stacking these mutations confers multiple CBIs resistance without compromising basal function of CSC complex or plant development.

**Fig. 6.**
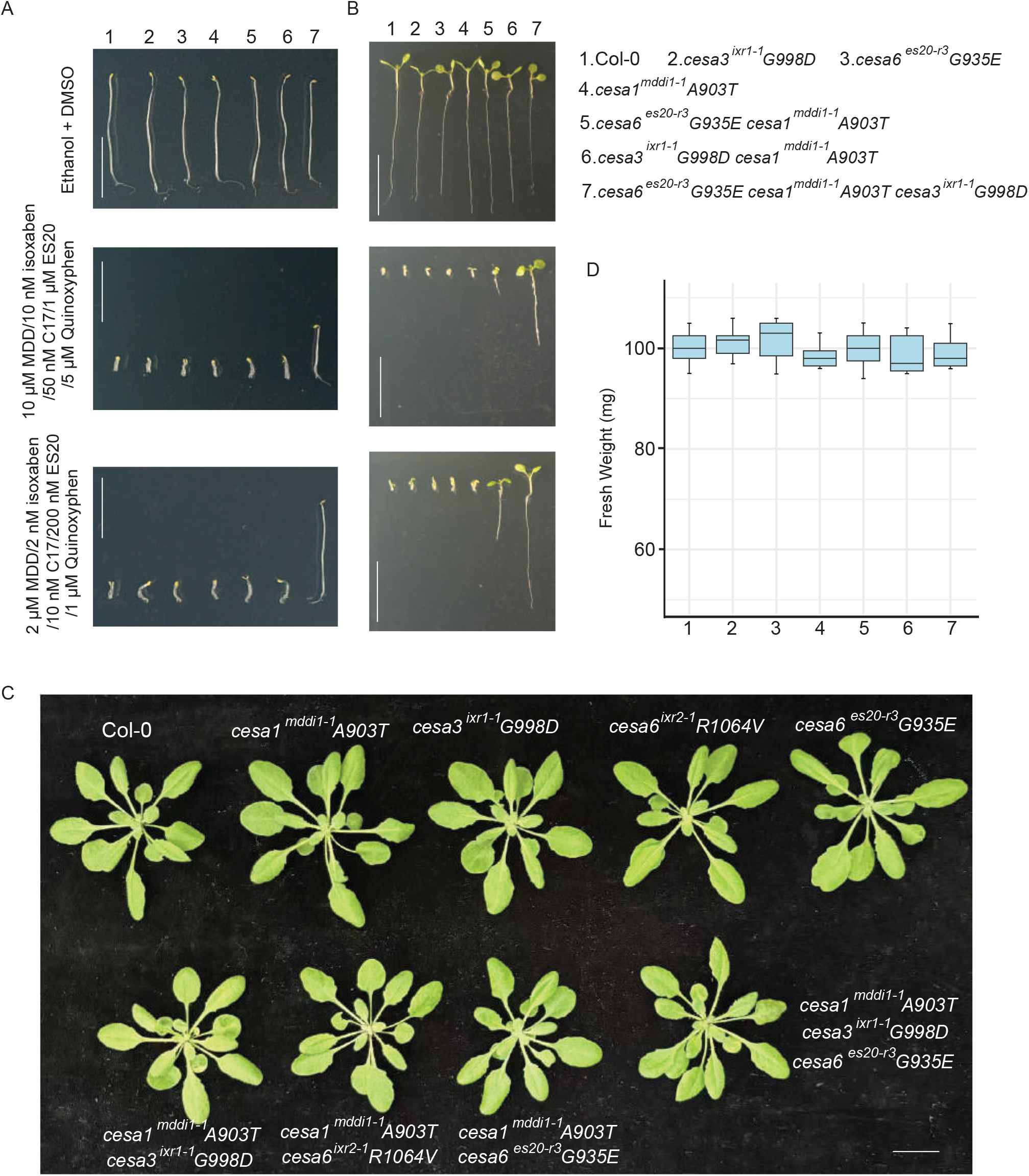
Stacking different mutations to generate multiple drug-resistant mutants. A.Representative image of Col-0, *cesa3*^*ixr1-1*^ *G998D, cesa6* ^*es20-r3*^ *G935E, cesa1*^*mddi1-1*^ *A903T, cesa3*^*ixr1-1*^ *G998D cesa1*^*mddi1-1*^ *A903T, cesa6* ^*es20-r3*^ *G935E cesa1*^*mddi1-1*^ *A903T* and *cesa6* ^*es20-r3*^ *G935E cesa1*^*mddi1-1*^ *A903T cesa3*^*ixr1-1*^ *G998D* hypocotyl growth of 5-day-old dark-grown Arabidopsis seedlings grown on ½ MS medium supplemented with low amount of Ethanol/DMSO, a combination treatment with 10 μM MDD/10 nM isoxaben/50 nM C17/1 μM ES20/5 μM Quinoxyphen, or a combination treatment with 2 μM MDD/2 nM isoxaben/10 nM C17/200 nM ES20/1 μM Quinoxyphen. Bars = 1.0 cm. B.Representative image of Col-0, *cesa3*^*ixr1-1*^ *G998D, cesa6* ^*es20-r3*^ *G935E, cesa1*^*mddi1-1*^ *A903T, cesa3*^*ixr1-1*^ *G998D cesa1*^*mddi1-1*^ *A903T, cesa6* ^*es20-r3*^ *G935E cesa1*^*mddi1-1*^ *A903T* and *cesa6* ^*es20-r3*^ *G935E cesa1*^*mddi1-1*^ *A903T cesa3*^*ixr1-1*^ *G998D* root growth of 7-day-old light-grown Arabidopsis seedlings grown on ½ MS medium supplemented with low amount of Ethanol/DMSO, a combination treatment with 10 μM MDD/10 nM isoxaben/50 nM C17/1 μM ES20/5 μM Quinoxyphen, or a combination treatment with 2 μM MDD/2 nM isoxaben/10 nM C17/200 nM ES20/1 μM Quinoxyphen. Bars = 1.0 cm. C.Representative image of 4-week-old of Col-0, *cesa1*^*mddi1-1*^ *A903T, cesa3*^*ixr1-1*^ *G998D, cesa6*^*ixr2-1*^ *R1064V, cesa6* ^*es20-r3*^ *G935E, cesa1*^*mddi1-1*^ *A903T cesa3*^*ixr1-1*^ *G998D, cesa1*^*mddi1-1*^ *A903T cesa6*^*ixr2-1*^ *R1064V, cesa1*^*mddi1-1*^ *A903T cesa6* ^*es20-r3*^ *G935E*, and *cesa1*^*mddi1-1*^ *A903T cesa3*^*ixr1-1*^ *G998D cesa6* ^*es20-r3*^ *G935E* Arabidopsis plants. Bars = 1.0 cm. D.Fresh weight of 4-week-old of Col-0, *cesa3*^*ixr1-1*^ *G998D, cesa6* ^*es20-r3*^ *G935E, cesa1*^*mddi1-1*^ *A903T, cesa3*^*ixr1-1*^ *G998D cesa1*^*mddi1-1*^ *A903T, cesa6* ^*es20-r3*^ *G935E cesa1*^*mddi1-1*^ *A903T* and *cesa6* ^*es20-r3*^ *G935E cesa1*^*mddi1-1*^ *A903T cesa3*^*ixr1-1*^ *G998D* Arabidopsis plants. Bars = 1.0 cm.

## Discussion

Bioactive natural products have long served as valuable resources for discovering potent herbicidal agents. CBIs are of particular interest as herbicide due to their ability to disrupt the formation of plant cell walls, thereby impairing plant growth. In this study, we identified MDD as a novel, natural product-derived CBI that inhibits plant cellulose synthesis by likely targeting the transmembrane domains of CESA1. MDD suppresses plant growth in both dicot and monocot species, demonstrating broad-spectrum applicability. Furthermore, the generation of multiple-drugs resistant mutants through stacking mutations in CESA subunits showcased a promising strategy for improving crop resistance to CBIs while informing fundamental mechanisms of cellulose biosynthesis.

Although most characterized CBIs are synthetic, a small but growing set of natural products also target cellulose biosynthesis. Thaxtomin A, produced by the plant-pathogenic bacterium *Streptomyces scabies*, is the best⍰characterized natural CBI. Thaxtomin A functions as a phytotoxin in plant-microbe interactions and is required for its pathogenicity for causing potato scab disease (38, 39). Applied to plants, Thaxtomin A disrupts cellulose synthesis in expanding plant cells, leading to characteristic cell swelling and root growth defects (40). Acetobixan is another natural CBI, which is isolated from a *Panicum virgatum* bacterial endophyte. Acetobixan induces radial cell swelling and growth inhibition in etiolated *Arabidopsis* seedlings, and triggers internalization of CESA6 in a manner reminiscent of isoxaben. Notably, acetobixan appears to act by a mechanism distinct from isoxaben and quinoxyphen because known mutants resistant to those CBIs do not show cross⍰resistance to acetobixan, and no acetobixan⍰resistant mutant has yet been identified (41). MDD joins this emerging class of natural CBIs and it is produced by *Aspergillus oryzae* and *Hamigera fusca*, with homologous BGCs conserved across various fungi species (30, 31). Further exploration of the role of MDD in these fungi, particularly in plant-microbe interactions, could provide valuable insights into its ecological function. Structurally, MDD differs significantly from synthetic CBIs like isoxaben, quinoxyphen, or C17 and natural CBIs like thaxtomin A, yet it exerts similar phenotypic effects, including inhibition of cellulose deposition and perturbation of cell wall integrity disruption. While the genetic cross⍰resistance patterns reported here suggest that MDD, quinoxyphen and C17 may perturb CSC function in related ways, definitive assignment of binding sites and modes requires direct biochemical and structural demonstration. Therefore, future work employing complementary approaches—quantitative binding assays, target⍰engagement methods, and structural studies of CESA domains or reconstituted CSC subcomplexes will be critical to pinpoint compound binding sites, discriminate primary versus indirect effects, and enable rational design of more potent and resistance⍰resilient CBIs.

The observation that MDD depletes CSC complexes from the plasma membrane suggests that it might interferes with their normal delivery or maintenance, potentially by disrupting CSC trafficking or destabilizing the complexes themselves. However, the absence of resistance to MDD in mutants lacking key vesicle trafficking regulators—such as CSI1, PATROL1, and SHOU4—indicates that the quantity of CSCs at the plasma membrane is not the sole factor affecting sensitivity to MDD. It is possible that MDD alters membrane composition or integrity, or may influence other, yet unidentified, trafficking or anchoring processes. Interestingly, MDD, quinoxyphen, and C17 insensitive mutants exhibit a degree of cross-resistance, but not others, suggesting that some substitution sites in their insensitive mutants may serve as herbicide binding sites. However, due to the structural diversity of these herbicides, not all substitutions for one herbicide necessarily behave the same way for another. As a result, these herbicides may disrupt the function or efficiency of CSCs, even while significantly affecting their abundance at the plasma membrane. This understanding emphasizes the complexity of the action of CBIs and highlights the need for further exploration into how MDD and other CBIs affects CSC functionality beyond simply altering their quantity at the plasma membrane. In addition, only two MDD⍰insensitive mutants were recovered from our EMS screen, while two additional C17⍰insensitive alleles showed partial resistance to MDD. These findings indicate that additional targeted CESA1 mutagenesis would be valuable to identify additional critical residues and refine the mode of action. Moreover, because the four residues implicated in MDD resistance are conserved across CESA family members, it will also be informative to test whether the corresponding substitutions in other CESA isoforms confer similar insensitivity.

Our transcriptomic and cell⍰biological data indicate that MDD impairs both cell expansion and proliferation. Short⍰term cell⍰wall perturbation by CBIs (for example, isoxaben) can activate cell⍰wall integrity (CWI) sensing and pattern⍰triggered immunity (PTI)⍰like transcriptional programs, accompanied by downregulation of growth-related genes (42). In our experiments, continuous MDD exposure led to downregulation of mitotic⍰ and cytokinesis⍰related genes. A parsimonious interpretation is that primary inhibition of cellulose biosynthesis perturbs cell⍰wall mechanics and integrity, which then feeds back on plant immunity, cell⍰cycle control and cytokinesis, producing fewer but swollen cells. The fact that point substitutions in CESA1 confer robust resistance strongly supports CESA1 as a principal functional target of MDD. Nevertheless, we cannot exclude additional targets that may contribute to the observed transcriptional and cellular phenotypes.

The multiple drug-resistant mutants created here remained morphologically normal while exhibiting robust resistance to multiple CBIs—including isoxaben, ES20, MDD, quinoxyphen, and C17—at concentrations that severely inhibit WT plants. From a weed management perspective, engineering crops with resistance to multiple CBIs is highly advantageous. Given the lack of cross-resistance between different inhibitor-resistant mutants, employing rotations or combinations of CBIs with partially overlapping but distinct targets sites can disperse selection pressure and substantially reduces the likelihood that weeds will acquire simultaneous resistance to all agents. Although the A903 residue affects resistance to multiple compounds, inclusion of additional herbicides in management strategies may further lowers the risk of broad⍰spectrum resistance. Therefore, expanding the range of available herbicides—particularly those that target distinct residues within cellulose synthases—offers greater flexibility and sustainability in integrated weed management programs.

## Materials and Methods

### Plant materials and growth conditions

All *Arabidopsis* plants used in this paper are Col-0 ecotype. *N. benthamiana, S. lycopersicum, Z. mays* and *Arabidopsis* plants were grown under standard condition with 16□h light/8□h dark at 22□°C (43). The CBI resistant lines used in this study were reported previously (13, 17, 32). The T-DNA insertion lines used in this study are *cesa1* heterozygous mutant (CS812877; *SAIL_278_E08*), *csi1-4* (44), *patrol1-2* (44) and *shou4-3*(37). Agrobacterium (AGL0 strain)-mediated floral dipping was used to generate all the transgenic plants. *pCESA1:YFP-CESA1, pCESA1:YFP-CESA1A903T* and *pCESA1:YFP-CESA1H024Y* constructs were transformed into *cesa1* heterozygous mutant. 16 transgenic T1 plants were selected on ½ MS medium containing hygromycin. Two T2 homozygous transgenic plants were tested for their sensitivity on ½ MS plates with 10 μM MDD.

### Compound isolation

Compound isolation was described previously (30). Briefly, the spores of *A. nidulans* transformed with corresponding plasmids were inoculated into 4 L CD-ST agar media and grown for 4 days at 28°C. The culture was extracted with 2 L ethyl acetate for two times and the solvent was evaporated to dryness under vacuum to obtain the crude extract. The crude extract was applied to normal-phase silica flash chromatography on a CombiFlash instrument with a gradient of ethyl acetate-hexanes (15:85 to 95:5). The fractions containing MDD were concentrated and subjected to reverse-phase C18 flash chromatography on a CombiFlash instrument with a gradient of CH_3_CN-H_2_O (40:60 to 60:40, 0.1% FA) to yield 31 mg of MDD (7.75 mg/L).

### RNA-seq

One week old of Col-0 seedlings treated with ethanol or MDD treatment were collected and ground into a fine powder with liquid nitrogen. Total RNA was extracted using the Direct-zol RNA Miniprep kit (Zymo) according to the manufacturer’s instructions. One microgram of total RNA was used to prepare the libraries for RNA sequencing (RNA-seq) following the TruSeq Stranded mRNA kit (Illumina), and the libraries were sequenced on a NovaSeq X Plus instruments (45).

### EMS Mutagenesis, mutant Screening and mapping

To obtain a mutagenized Arabidopsis population, approximately 5,000 Col-0 seeds were mutagenized with ethyl methanesulfonate (EMS) for 14□h. Roughly 30,000 M2 plants representing ∼2,000 M1 families were grown on ½ MS medium with 10 μM MDD. To map *mddi* mutants, homozygous *mddi* mutants were backcrossed with Col-0 to generate a mapping population. Sixty F2 plants insensitive to MDD were selected for Illumina sequencing. Single nucleotide polymorphism (SNP) frequency was calculated.

### Plasmid construction

For *pCESA1:YFP-CESA1, pCESA1:YFP-CESA1A903T* and *pCESA1:YFP-CESA1H024Y*, the genomic DNA sequences were amplified from Col-0, homozygous *mddi1-1* and *mddi1-2* plants respectively and cloned into pENTR/D-TOPO vectors (Invitrogen). Inserts were then transferred to the destination vector pEG302-GW by LR reaction (LR Clonase II, Invitrogen). Primers used in this study are presented in Supplementary Table 1.

### Structural Modeling of the CESA1

The general topology of CESA6 was predicted using the UniProt (https://www.uniprot.org) and PredictProtein (https://www.predictprotein.org) servers, and the cartoon was drawn using the Protter program (http://wlab.ethz.ch/protter/start/).

### Crystalline Cellulose Content Measurement

WT and *mddi Arabidopsis* seeds were sown on ½ MS plates with ethanol or 10 μM MDD for 7 days. Seedlings were washed with double-distilled water three times to remove seed coats and any residue from the growth medium and then ground into a fine powder in liquid nitrogen. The powder was extracted twice with 80% ethanol, once with 100% ethanol, once with 1:1 (v/v) methanol: CHCl3, and once with acetone. The resulting insoluble cell wall fraction was dried in a fume hood for 2 d and weighed. Cellulose content was measured by the Updegraff method (32). Briefly, the cell wall material was hydrolyzed with Updegraff reagent (acetic acid: nitric acid: water, 8:1:2 [v/v/v]) to yield crystalline cellulose. The residual pellet obtained after the hydrolysis was rinsed twice with 10 volumes of water to remove acetic and nitric acids. Air drying of the pellet was used to remove excess water. Crystalline cellulose was hydrolyzed to Glc using 72% (v/v) sulfuric acid. Glc concentration was measured via a colorimetric method by developing color in Anthrone reagent (freshly prepared 2 mg/mL anthrone in concentrated sulfuric acid) and reading OD625 plate reader (12).

### Live-cell imaging with spinning-disk confocal microscopy

Confocal microscopy experiments were performed using the LSM 980 confocal microscope or Evident FV4000 confocal microscopy. The root samples were collected from 1-week-old seedlings grown on ½ MS plates with ethanol or 10 μM MDD under standard condition with 16□h light/8□h dark at 22□°C. The hypocotyl samples were collected from 4-day-old seedlings grown on ½ MS plates with ethanol or 10 μM MDD under dark condition. Propidium iodide was used to stain cells. To examine the localization of the CSCs at the plasma membrane, the seedlings of GFP-CESA3 grown on ½ MS plates for 4 days were treated with ethanol or 100 μM MDD for 5 hr. GFP fluorescence was excited with a 488-nm laser and the emissions were collected using a 500-nm to 580-nm laser filter. For time series observation of GFP-CESA3, the pictures were taken every 10 s per picture for 4 min in total.

### Image analysis

Image analysis was performed using Fiji/ImageJ software. For CESA particle dynamic analyses, 5-min time-lapse series with 5-s intervals were collected. Average intensity projections were generated to identify the trajectories of the CSC particles. Image drift was corrected by the StackReg plugin. The moving rate of CESA3 was qualified by using Kymograph analysis as described (Bringmann et al. 2012). In brief, after each group of frames were stacked, kymographs were made using the multiple kymograph plugin for ImageJ (http://www.embl.de/eamnet/html/body_kymograph.html), and then, particle velocities were calculated from the slopes of kymographs.

## Acknowledgments

We thank Dr. Charles T. Anderson at The Pennsylvania State University for sharing the GFP-CESA3; mCherry-TUA5 seeds, Dr. Lieven De Veylder at Ghent University for the C17 resistant EMS mutants, and Dr. Christopher J. Staiger for the ES20 resistant EMS mutants. We also thank Mahnaz Akhavan, Suhua Feng and the UCLA BSCRC BioSequencing Core for sequencing support. This work was supported from Agricultural Science and Technology Major Project and by “the Fundamental Research Funds for the Central Universities”+ 226-2025-00083 to Z.W. and funded by NIFA (2021-67013-34259) and NSF Award 2434499 to Y.T. and S.E.J.. S.E.J. is an Investigator of the Howard Hughes Medical Institute.

## Data and materials availability

All the high-throughput sequencing data generated in this study is accessible at the Genome Sequence Archive database in National Genomics Data Center (https://ngdc.cncb.ac.cn/) under the accession PRJCA055487.

## References

1. P. Purushotham, R. Ho, J. Zimmer, Architecture of a catalytically active homotrimeric plant cellulose synthase complex. Science 369, 1089–1094 (2020).

2. J. Du et al., Evidence for Plant-Conserved Region Mediated Trimeric CESAs in Plant Cellulose Synthase Complexes. Biomacromolecules 23, 3663–3677 (2022).

3. J. L. Morgan, J. Strumillo, J. Zimmer, Crystallographic snapshot of cellulose synthesis and membrane translocation. Nature 493, 181–186 (2013).

4. X. Zhang et al., Structural insights into homotrimeric assembly of cellulose synthase CesA7 from Gossypium hirsutum. Plant Biotechnol J 19, 1579–1587 (2021).

5. N. G. Taylor, R. M. Howells, A. K. Huttly, K. Vickers, S. R. Turner, Interactions among three distinct CesA proteins essential for cellulose synthesis. Proc Natl Acad Sci U S A 100, 1450–1455 (2003).

6. T. Desprez et al., Organization of cellulose synthase complexes involved in primary cell wall synthesis in Arabidopsis thaliana. Proc Natl Acad Sci U S A 104, 15572–15577 (2007).

7. S. Persson et al., Genetic evidence for three unique components in primary cell-wall cellulose synthase complexes in Arabidopsis. Proc Natl Acad Sci U S A 104, 15566–15571 (2007).

8. J. K. Polko, J. J. Kieber, The Regulation of Cellulose Biosynthesis in Plants. Plant Cell 31, 282–296 (2019).

9. R. T. Larson, H. E. McFarlane, Small but Mighty: An Update on Small Molecule Plant Cellulose Biosynthesis Inhibitors. Plant Cell Physiol 62, 1828–1838 (2021).

10. J. M. Alvarez, Cellulose biosynthesis inhibitors as tools for research of cell wall structural plasticity, Cell biology research progress (Nova Science Publishers, Hauppauge, N.Y, 2012).

11. C. Brabham, S. Debolt, Chemical genetics to examine cellulose biosynthesis. Front Plant Sci 3, 309 (2012).

12. L. Huang et al., Endosidin20 Targets the Cellulose Synthase Catalytic Domain to Inhibit Cellulose Biosynthesis. Plant Cell 32, 2141–2157 (2020).

13. Z. Hu et al., Mitochondrial Defects Confer Tolerance against Cellulose Deficiency. Plant Cell 28, 2276–2290 (2016).

14. I. Shim et al., Alleles Causing Resistance to Isoxaben and Flupoxam Highlight the Significance of Transmembrane Domains for CESA Protein Function. Front Plant Sci 9, 1152 (2018).

15. S. DeBolt et al., Morlin, an inhibitor of cortical microtubule dynamics and cellulose synthase movement. Proc Natl Acad Sci U S A 104, 5854–5859 (2007).

16. M. D. Lazzaro, J. M. Donohue, F. M. Soodavar, Disruption of cellulose synthesis by isoxaben causes tip swelling and disorganizes cortical microtubules in elongating conifer pollen tubes. Protoplasma 220, 201–207 (2003).

17. W. R. Scheible, R. Eshed, T. Richmond, D. Delmer, C. Somerville, Modifications of cellulose synthase confer resistance to isoxaben and thiazolidinone herbicides in Arabidopsis Ixr1 mutants. Proc Natl Acad Sci U S A 98, 10079–10084 (2001).

18. D. M. Harris et al., Cellulose microfibril crystallinity is reduced by mutating C-terminal transmembrane region residues CESA1A903V and CESA3T942I of cellulose synthase. Proc Natl Acad Sci U S A 109, 4098–4103 (2012).

19. C. Brabham et al., Indaziflam herbicidal action: a potent cellulose biosynthesis inhibitor. Plant Physiol 166, 1177–1185 (2014).

20. D. R. Heim, I. M. Larrinua, M. G. Murdoch, J. L. Roberts, Triazofenamide is a cellulose biosynthesis inhibitor. Pestic Biochem Phys 59, 163–168 (1998).

21. A. Cano-Delgado, S. Penfield, C. Smith, M. Catley, M. Bevan, Reduced cellulose synthesis invokes lignification and defense responses in Arabidopsis thaliana. Plant J 34, 351–362 (2003).

22. I. Sorensen et al., The charophycean green algae provide insights into the early origins of plant cell walls. Plant J 68, 201–211 (2011).

23. A. R. Paredez, C. R. Somerville, D. W. Ehrhardt, Visualization of cellulose synthase demonstrates functional association with microtubules. Science 312, 1491–1495 (2006).

24. Z. Hu et al., Genome Editing-Based Engineering of CESA3 Dual Cellulose-Inhibitor-Resistant Plants. Plant Physiol 180, 827–836 (2019).

25. L. Huang, C. Zhang, The Mode of Action of Endosidin20 Differs from That of Other Cellulose Biosynthesis Inhibitors. Plant Cell Physiol 61, 2139–2152 (2021).

26. L. Sethaphong et al., Tertiary model of a plant cellulose synthase. Proc Natl Acad Sci U S A 110, 7512–7517 (2013).

27. P. J. Rutledge, G. L. Challis, Discovery of microbial natural products by activation of silent biosynthetic gene clusters. Nat Rev Microbiol 13, 509–523 (2015).

28. N. Ziemert, M. Alanjary, T. Weber, The evolution of genome mining in microbes - a review. Nat Prod Rep 33, 988–1005 (2016).

29. P. Cimermancic et al., Insights into Secondary Metabolism from a Global Analysis of Prokaryotic Biosynthetic Gene Clusters. Cell 158, 412–421 (2014).

30. M. Liu et al., AoiQ Catalyzes Geminal Dichlorination of 1,3-Diketone Natural Products. J Am Chem Soc 143, 7267–7271 (2021).

31. C. Almeida et al., Non-geminal Aliphatic Dihalogenation Pattern in Dichlorinated Diaporthins from Hamigera fusca NRRL 35721. J Nat Prod 81, 1488–1492 (2018).

32. T. Desprez et al., Resistance against herbicide isoxaben and cellulose deficiency caused by distinct mutations in same cellulose synthase isoform CESA6. Plant Physiol 128, 482–490 (2002).

33. N. Hoffmann, S. King, A. L. Samuels, H. E. McFarlane, Subcellular coordination of plant cell wall synthesis. Dev Cell 56, 933–948 (2021).

34. M. Bringmann et al., POM-POM2/CELLULOSE SYNTHASE INTERACTING1 Is Essential for the Functional Association of Cellulose Synthase and Microtubules in. Plant Cell 24, 163–177 (2012).

35. S. D. Li, L. Lei, C. R. Somerville, Y. Gu, Cellulose synthase interactive protein 1 (CSI1) links microtubules and cellulose synthase complexes. P Natl Acad Sci USA 109, 185–190 (2012).

36. X. Y. Zhu, S. D. Li, S. Q. Pan, X. R. Xin, Y. Gu, CSI1, PATROL1, and exocyst complex cooperate in delivery of cellulose synthase complexes to the plasma membrane (vol 115, pg E3578, 2018). P Natl Acad Sci USA 115, E5635–E5635 (2018).

37. J. K. Polko et al., SHOU4 Proteins Regulate Trafficking of Cellulose Synthase Complexes to the Plasma Membrane. Curr Biol 28, 3174–3182 e3176 (2018).

38. C. Goyer, J. Vachon, C. Beaulieu, Pathogenicity of Streptomyces scabies Mutants Altered in Thaxtomin A Production. Phytopathology 88, 442–445 (1998).

39. M. V. Joshi et al., The AraC/XylS regulator TxtR modulates thaxtomin biosynthesis and virulence in Streptomyces scabies. Mol Microbiol 66, 633–642 (2007).

40. W. R. Scheible et al., An Arabidopsis mutant resistant to thaxtomin A, a cellulose synthesis inhibitor from Streptomyces species. Plant Cell 15, 1781–1794 (2003).

41. Y. Xia et al., Acetobixan, an inhibitor of cellulose synthesis identified by microbial bioprospecting. Plos One 9, e95245 (2014).

42. K. Zhai, J. Rhodes, C. Zipfel, A peptide-receptor module links cell wall integrity sensing to pattern-triggered immunity. Nat Plants 10, 2027–2037 (2024).

43. Z. Wu et al., REM transcription factors and GDE1 shape the DNA methylation landscape through the recruitment of RNA polymerase IV transcription complexes. Nat Cell Biol 27, 1136–1147 (2025).

44. L. Liu et al., Actomyosin and CSI1/POM2 cooperate to deliver cellulose synthase from Golgi to cortical microtubules in Arabidopsis. Nat Commun 14, 7442 (2023).

45. M. Wang et al., Arabidopsis TRB proteins function in H3K4me3 demethylation by recruiting JMJ14. Nat Commun 14, 1736 (2023).

