## Supplemental Figure 1-9 for "A Fungal Natural Product that Inhibits Plant Cellulose Biosynthesis by disrupting Cellulose Synthase Complexes"

**This PDF file includes:**

Figures S1 to S9
Tables S1

Fig S1

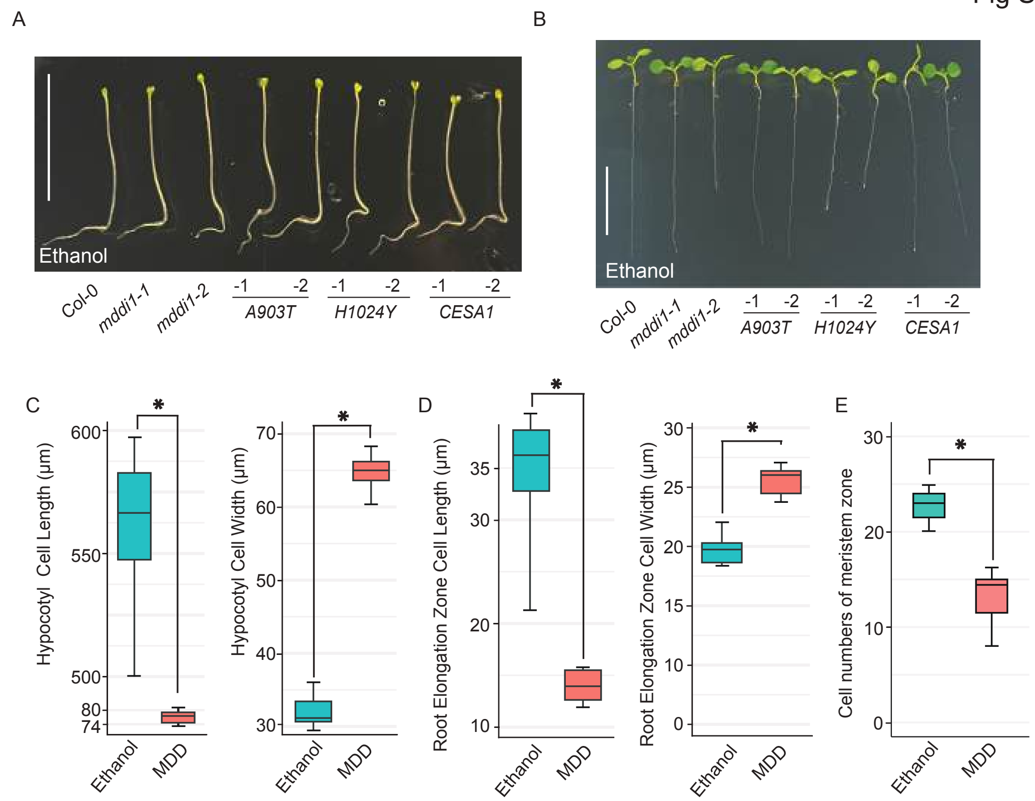


**Fig. S1. MDD induces swelling in Arabidopsis hypocotyl and root cells.**
A. Representative image of hypocotyl cells of 5-day-old dark-grown Arabidopsis seedlings grown on ½ MS medium supplemented with ethanol. Bars = 1.0 cm.
B. Representative image of root cells of 7-day-old light-grown Arabidopsis seedlings grown on ½ MS medium supplemented with ethanol. Bars = 1.0 cm.
C. Length and width of hypocotyl cells of 4-day-old dark-grown Arabidopsis seedlings grown on ½ MS medium supplemented with ethanol or 10 μM MDD. Statistical significance is indicated by different letters ($p < 0.01$). Error bars represent means ± standard deviations (n = 3).
D. Length and width of elongation zone root cells of 4-day-old light-grown Arabidopsis seedlings grown on ½ MS medium supplemented with ethanol or 10 μM MDD. Statistical significance is indicated by different letters ($p < 0.01$). Error bars represent means ± standard deviations (n = 3).
E. Cell numbers of meristem zone root cells of 4-day-old light-grown Arabidopsis seedlings grown on ½ MS medium supplemented with ethanol or 10 μM MDD. Statistical significance is indicated by different letters ($p < 0.01$). Error bars represent means ± standard deviations (n = 3).

Fig S2

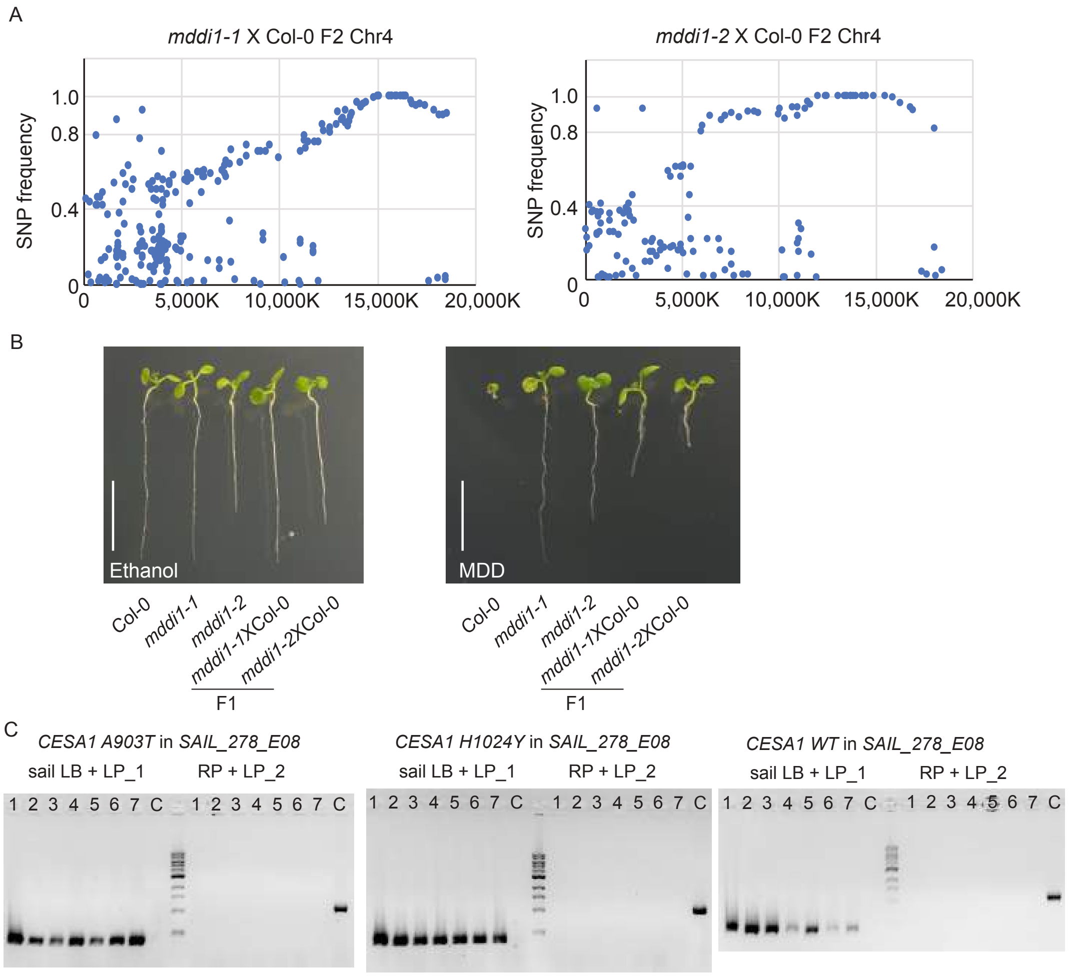


**Fig. S2. Resistance to MDD is conferred by semi-dominant CESA1 alleles.**
A. Dot plots of SNP frequency on chromosome 4 in *mddi1-1, mddi1-2* bulked backcross segregants.
B. Representative root image of *mddi1-1* and *mddi1-2* F1 backcrossed Arabidopsis seedlings grown on ½ MS medium supplemented with ethanol or 10 μM MDD. Bars = 1.0 cm.
C. Genotyping result showing the identification of *cesa1* null mutant when N-terminal GFP-tagged CESA1 A903T or CESA1 H1024Y or WT CESA1 were transformed into heterozygous *cesa1* knock-out mutants. The presence of T-DNA insertion in all T3 progenies from selected T2 lines confirmed homozygosity of the T-DNA insertion.

Fig S3

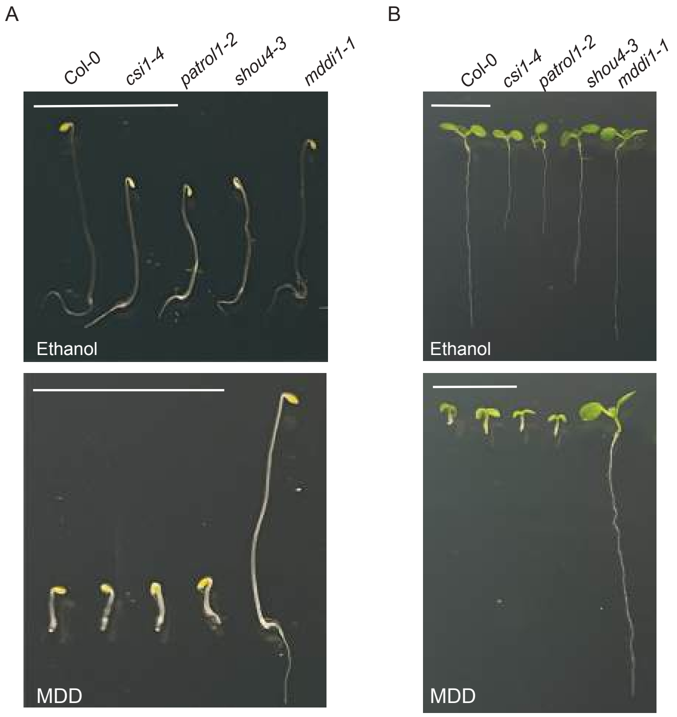


**Fig. S3. MDD sensitivity remains unaltered by CSC complex quantity or regulatory mechanisms at the plasma membrane.**

A. Representative image of 5-day-old dark-grown indicated Arabidopsis seedlings grown on ½ MS medium supplemented with ethanol or 10 μM MDD. Bars = 1.0 cm.

B. Representative image of 7-day-old light-grown indicated Arabidopsis seedlings grown on ½ MS medium supplemented with ethanol or 10 μM MDD. Bars = 1.0 cm.

Fig S4

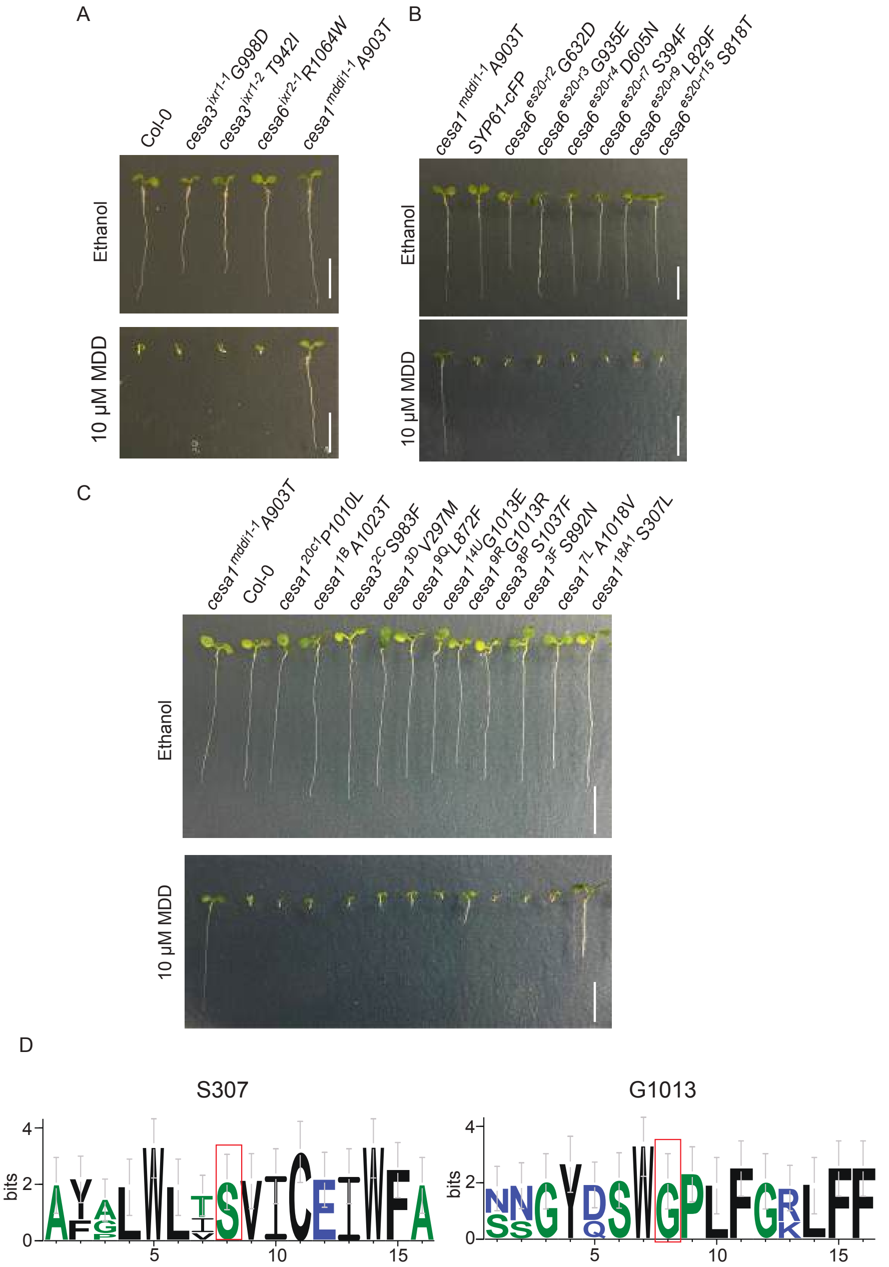


**Fig. S4. Isoxaben, ES20-resistant mutants are susceptible to MDD.**

A. Representative image of *cesa3*$^{ixr1\text{-}1}$ *G998D*, *cesa3*$^{ixr1\text{-}2}$ *T942I*, *cesa6*$^{ixr2\text{-}1}$ *R1064W* and *cesa1*$^{mddi1\text{-}1}$ *A903T* root growth of 7-day-old light-grown Arabidopsis seedlings grown on ½ MS medium supplemented with ethanol or 10 µM MDD. Bars = 1.0 cm.

B. Representative image of *cesa1*$^{mddi1\text{-}1}$ *A903T*, SYP61-CFP, *cesa6*$^{es20\text{-}r2}$ *G632D*, *cesa6*$^{es20\text{-}r3}$ *G935E*, *cesa6*$^{es20\text{-}r4}$ *D605N*, *cesa6*$^{es20\text{-}r7}$ *S394F*, *cesa6*$^{es20\text{-}r9}$ *L829F*, and *cesa6*$^{es20\text{-}r15}$ *S818T* root growth of 7-day-old light-grown Arabidopsis seedlings grown on ½ MS medium supplemented with ethanol or 10 µM MDD. Bars = 1.0 cm.

C. Representative image of *cesa1*$^{mddi1\text{-}1}$ *A903T*, Col-0, *cesa1*$^{20c1}$ *P1010L*, *cesa1*$^{1B}$ *A1023T*, *cesa3*$^{2C}$ *S983F*, *cesa1*$^{3D}$ *V297M*, *cesa1*$^{9Q}$ *L872F*, *cesa1*$^{14U}$ *G1013E*, *cesa1*$^{9R}$ *G1013R*, *cesa3*$^{8P}$ *S1037F*, *cesa1*$^{3F}$ *S892N*, *cesa1*$^{7L}$ *A1018V*, and *cesa1*$^{18A1}$ *S307L* root growth of 7-day-old light-grown Arabidopsis seedlings grown on ½ MS medium supplemented with ethanol or 10 µM MDD. Bars = 1.0 cm.

D. Sequence logo assessment of residues in the additional mutation regions of primary cell wall CESA proteins illustrates the location and conservation of the mutated serine and glycine residues in CESA1.

Fig S5

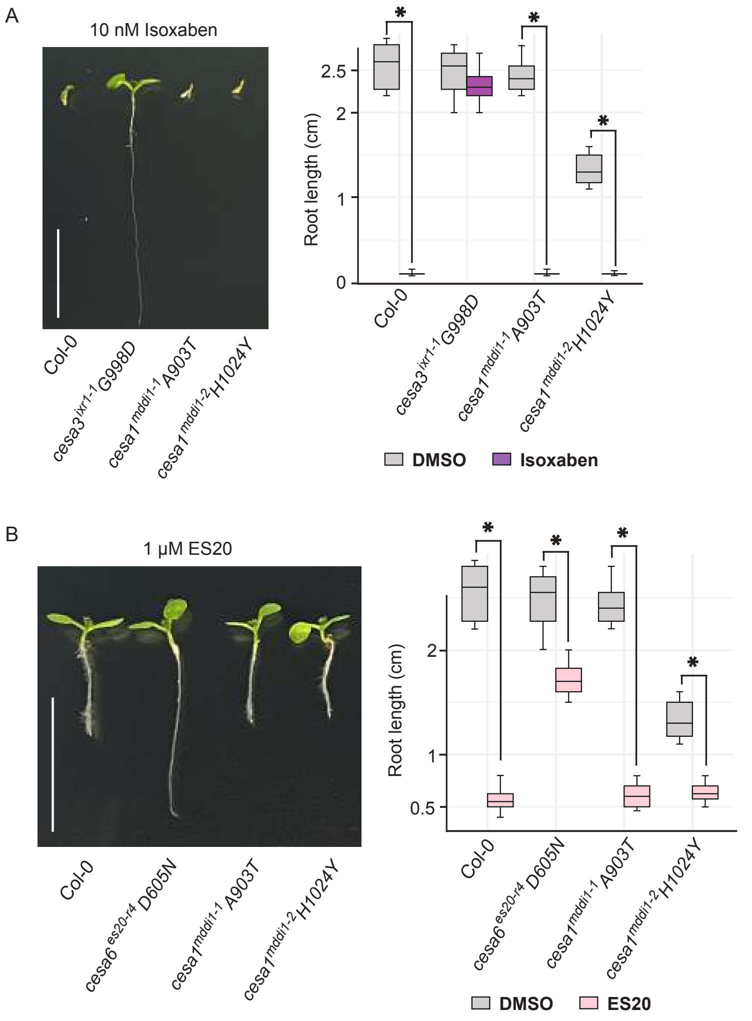


**Fig. S5. *mddi* mutants are sensitive to isoxaben and ES20.**

A. Representative image and quantification of Col-0, *cesa3*$^{ixr1\text{-}1}$ *G998D*, *cesa1*$^{mddi1\text{-}1}$ *A903T* and *cesa1*$^{mddi1\text{-}2}$ *H1024Y* root growth of 7-day-old light-grown Arabidopsis seedlings grown on ½ MS medium supplemented with 10 nM Isoxaben. Bars = 1.0 cm.

B. Representative image and quantification of Col-0, *cesa6*$^{es20\text{-}r4}$ *D605N*, *cesa1*$^{mddi1\text{-}1}$ *A903T* and *cesa1*$^{mddi1\text{-}2}$ *H1024Y* root growth of 7-day-old light-grown Arabidopsis seedlings grown on ½ MS medium supplemented with 1 µM ES20. Bars = 1.0 cm.

Fig S6

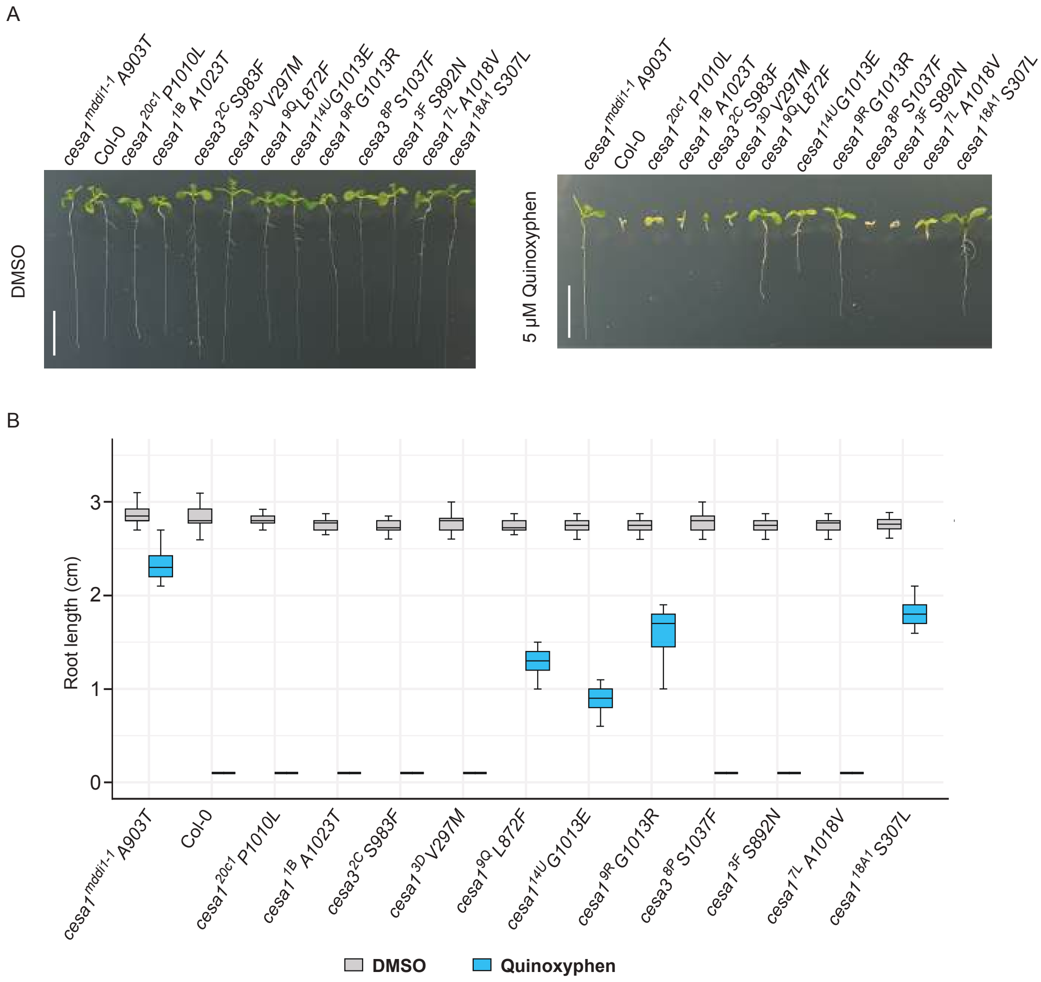


**Fig. S6. C17 insensitive mutants showed partial resistant to quinoxyphen.**
A-B. Representative image (A) and quantification (B) of *cesa1*$^{mddi1\text{-}1}$ *A903T*, Col-0, *cesa1* $^{20c1}$ *P1010L*, *cesa1* $^{1B}$ *A1023T*, *cesa3* $^{2C}$ *S983F*, *cesa1* $^{3D}$ *V297M*, *cesa1* $^{9Q}$ *L872F*, *cesa1* $^{14U}$ *G1013E*, *cesa1* $^{9R}$ *G1013R*, *cesa3* $^{8P}$ *S1037F*, *cesa1* $^{3F}$ *S892N*, *cesa1* $^{7L}$ *A1018V*, and *cesa1*$^{18A1}$ *S307L* root growth of 7-day-old light-grown Arabidopsis seedlings grown on ½ MS medium supplemented with DMSO or 5 µM Quinoxyphen. Bars = 1.0 cm.

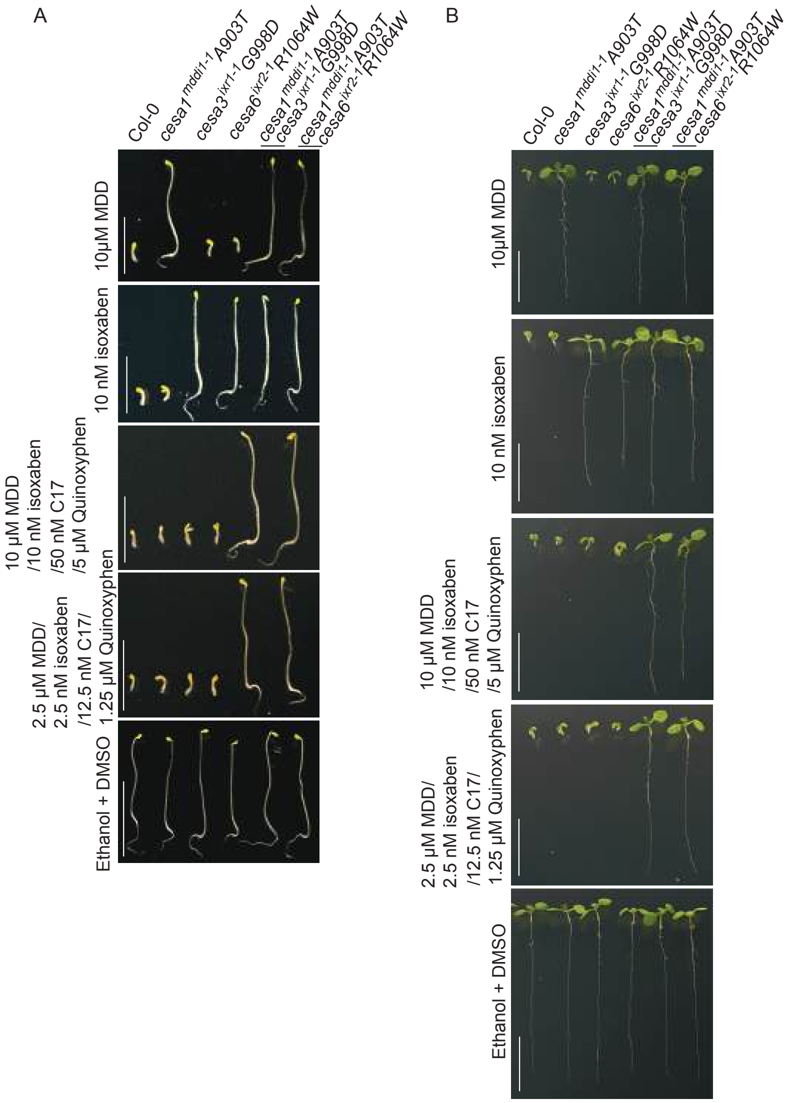


**Fig. S7. Generation of MDD/isoxaben/C17/Quinoxyphen resistant mutants.**
A. Representative image of Col-0, *cesa1*$^{mddi1\text{-}1}$ *A903T*, *cesa3*$^{ixr1\text{-}1}$ *G998D*, *cesa6*$^{ixr2\text{-}1}$ *R1064W*, *cesa1*$^{mddi1\text{-}1}$ *A903T cesa3*$^{ixr1\text{-}1}$ *G998D*, and *cesa1*$^{mddi1\text{-}1}$ *A903T cesa6*$^{ixr2\text{-}1}$ *R1064W* hypocotyl growth of 5-day-old dark-grown Arabidopsis seedlings grown on ½ MS medium supplemented with 10 μM MDD, 10 nM isoxaben, a combination treatment with 10 μM MDD/10 nM isoxaben/50 nM C17/5 μM Quinoxyphen, a combination treatment with 2.5 μM MDD/2.5 nM isoxaben/12.5 nM C17/1.25 μM Quinoxyphen. Bars = 1.0 cm.
B. Representative image of Col-0, *cesa1*$^{mddi1\text{-}1}$ *A903T*, *cesa3*$^{ixr1\text{-}1}$ *G998D*, *cesa6*$^{ixr2\text{-}1}$ *R1064W*, *cesa1*$^{mddi1\text{-}1}$ *A903T cesa3*$^{ixr1\text{-}1}$ *G998D*, and *cesa1*$^{mddi1\text{-}1}$ *A903T cesa6*$^{ixr2\text{-}1}$ *R1064W* root growth of 7-day-old light-grown Arabidopsis seedlings grown on ½ MS medium supplemented with 10 μM MDD, 10 nM isoxaben, a combination treatment with 10 μM MDD/10 nM isoxaben/50 nM C17/5 μM Quinoxyphen, or a combination treatment with 2.5 μM MDD/2.5 nM isoxaben/12.5 nM C17/1.25 μM Quinoxyphen. Bars = 1.0 cm.

Fig S8

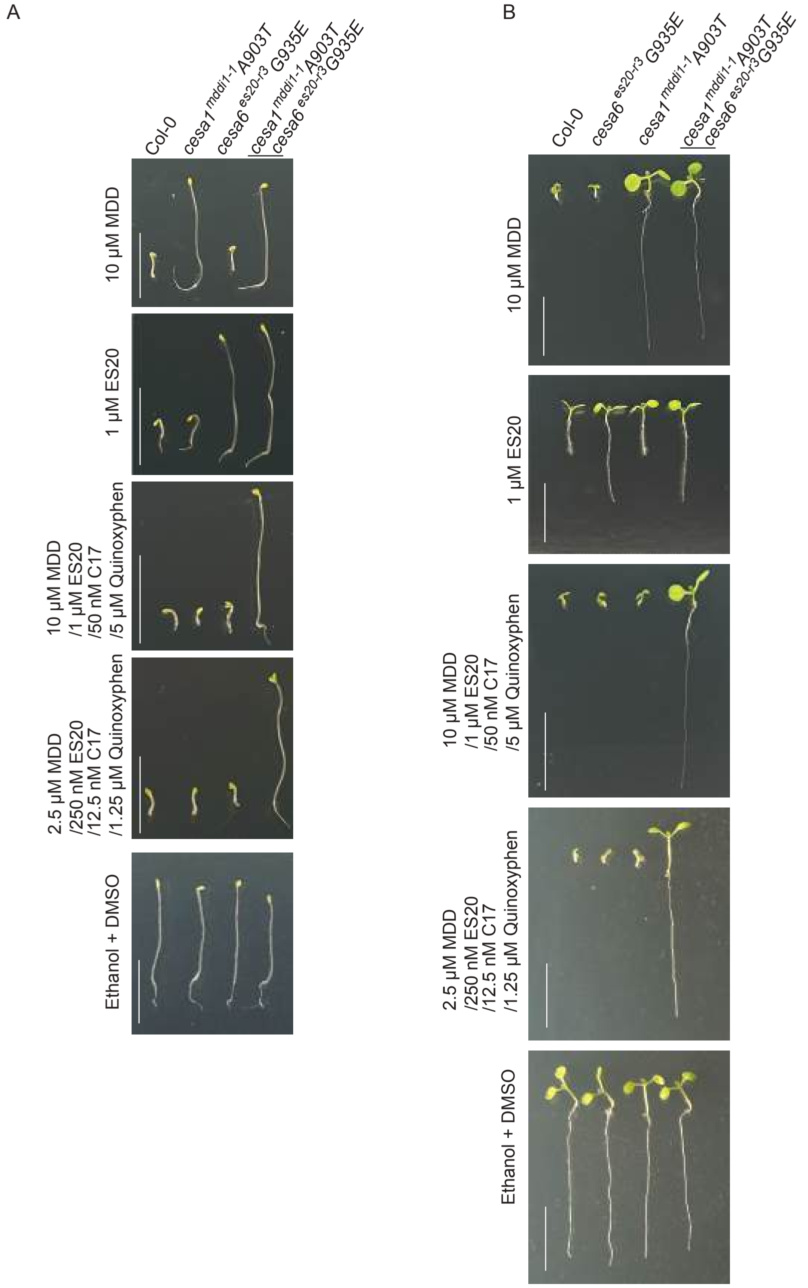


**Fig. S8. Generation of MDD/ES20/C17/Quinoxyphen resistant mutants.**
A. Representative image of Col-0, *cesa1*$^{mddi1-1}$ *A903T*, *cesa6* $^{es20-r3}$ *G935E*, and *cesa1*$^{mddi1-1}$ *A903T cesa6* $^{es20-r3}$ *G935E* hypocotyl growth of 5-day-old dark-grown Arabidopsis seedlings grown on ½ MS medium supplemented with 10 µM MDD, 1 µM ES20, a combination treatment with 10 µM MDD/1 µM ES20/50 nM C17/5 µM Quinoxyphen, a combination treatment with 2.5 µM MDD/250 nM ES20/12.5 nM C17/1.25 µM Quinoxyphen. Bars = 1.0 cm.
B. Representative image of Col-0, *cesa1*$^{mddi1-1}$ *A903T*, *cesa6* $^{es20-r3}$ *G935E*, and *cesa1*$^{mddi1-1}$ *A903T cesa6* $^{es20-r3}$ *G935E* root growth of 7-day-old light-grown Arabidopsis seedlings grown on ½ MS medium supplemented with 10 µM MDD, 1 µM ES20, a combination treatment with 10 µM

MDD/1 μM ES20/50 nM C17/5 μM Quinoxyphen, a combination treatment with 2.5 μM MDD/250 nM ES20/12.5 nM C17/1.25 μM Quinoxyphen. Bars = 1.0 cm.

**Supplemental Table1. Primers used in this study**

| Primer name | primer sequence |
|---|---|
| SAIL_278_E08_LP_1 | CCAGAACTGCTCGTTCCTCC |
| SAIL_278_E08_LP_2 | AGCTAATCTTGGACATGGAA |
| SAIL_278_E08 _RP | CCTGCGTTCAAGGGATCTGC |
| CESA1-F1 | CCGCGGCCGCCCCCTTCACCCGAAGTCGGATTCGACATCC |
| CESA1-R1 | CTCCTCGCCCTTGCTCACCATCGCAGCCACCGACACACAGA |
| CESA1-F2 | TCTGTGTGTCGGTGGCTGCGATGGTGAGCAAGGGCGAGGAG |
| CESA1-R2 | AAGCCGGCACTGGCCTCCATAGCCTTGTACAGCTCGTCCA |
| CESA1-F3 | TGGACGAGCTGTACAAGGCTATGGAGGCCAGTGCCGGCTT |
| CESA1-R3 | GGGTCGGCGCGCCCACCCTTCTAAAAGACACCTCCTTTGC |
| ixr1_F | AATGGTGGAGAAACGAGCAG |
| ixr1_R | CAACAGTTGATTCCACATTC |
| es20r3_F | TGCGAGTATCCTCTTCATGG |
| es20r3_R | GGGATGAGAAGTGAAGTCCA |
| ixr2_F | AGACTGTTCTTTGCACTTTG |
| ixr2_R | TCACAAGCAGTCTAAACCAC |
| mddi1-F | ACATTGTCCTATCTGGTATG |
| mddi1-R | AGTAGACCTTGGAAGACAGC |
